# Modeling cell-cell interactions to advance drug discovery in Idiopathic Pulmonary Fibrosis

**DOI:** 10.64898/2026.01.29.702646

**Authors:** Chandani Sen, Justin Langerman, Konstantinos Alysandratos, Andrea B Alber, Caroline Cherry, Kristen Castillo, Sarah Siegel, Rachana R Chandran, WooSuk Choi, Tammy Rickabaugh, Darrell N Kotton, Kathrin Plath, Brigitte N Gomperts

## Abstract

**Background:** Idiopathic Pulmonary Fibrosis (IPF) is characterized by scarring and remodeling of lung tissue, leading to progressive pulmonary dysfunction. Currently, very little is known about the steps involved in disease initiation and progression because models of IPF poorly replicate these processes. However, understanding the pathogenesis of IPF is essential for developing effective therapies. To address this, we have developed a scaffold-based co-culture IPF organoid that uses healthy and diseased human primary and iPSC-derived alveolar epithelial cells and fibroblasts to recapitulate the cellular communication and cell fate during progressive fibrosis.

**Methods:** We generated microbead scaffolds that mimic alveolar air-sacs and coated them in a rotating bioreactor with primary lung fibroblasts and induced pluripotent stem cell-derived type 2 alveolar cells (iAT2s). iAT2s with the surfactant protein C (SFTPC) I73T variant and the syngeneic corrected control iAT2s were cultured with primary healthy and IPF fibroblasts in different combinations. The epithelial-mesenchymal interactions during fibrosis initiation and progression were evaluated by single-cell RNA sequencing.

**Results:** We found that the interaction between epithelial cells and fibroblasts plays a key role in inducing fibrotic responses in this model, with the secretion of chemokines, cytokines, TGFβ, and matrix metalloproteinases that mirror those observed in the serum of patients with pulmonary fibrosis. Single-cell RNA sequencing revealed the emergence of many cell subtypes observed in progressive lung fibrosis, along with key cellular interactions that correlated with the initial upregulation of fibrosis pathways, extracellular matrix (ECM) remodeling, inflammation, and changes in lipid metabolism. The anti-fibrotic compounds, Nintedanib and the TGFβ inhibitor, SB431542, demonstrated dose-dependent efficacy in the model, with IC50 values comparable to those observed in the clinic, and significantly reduced secretion of fibrosis-related factors.

**Conclusion:** Overall, this study shows that the three-dimensional, reductionist, cell co-culture organoid effectively models several components of progressive lung fibrosis, facilitating the investigation of epithelial-mesenchymal interactions and serving as a patient-relevant model to better predict the efficacy of therapeutics in the clinic.

## Background

Idiopathic Pulmonary Fibrosis (IPF) is a chronic, progressive, and ultimately fatal interstitial lung disease characterized by excessive extracellular matrix (ECM) deposition, aberrant epithelial repair, and relentless fibrotic remodeling of the alveolar space [1]. The overall 5-year survival rate is only around 20%, and there is great variability in time to progression, making it very challenging to determine the prognosis for an individual patient. Despite significant advances in understanding the histopathological features of IPF, the underlying cause and precise cellular and molecular mechanisms of disease initiation and progression remain poorly defined, which continues to hinder the development of effective therapeutics [2]. There have been many failed clinical therapeutic trials for IPF [3, 4], and reasons for this include the complexity of the disease and poor understanding of its pathophysiology, the lack of models that truly recapitulate the progressive nature of the disease, and issues with identifying the correct efficacy endpoints in clinical trials. There is also a major need to identify biomarkers of IPF, particularly biomarkers of response to therapy [5]. The Prognostic Lung Fibrosis Consortium (PROLIFIC) identified five serum proteins (MMP7, TNC, sICAM1, SFTPD, and CXCL1) that predicted both lung function decline and transplant-free survival in one year, and these show promise for clinical utility.

Current FDA-approved treatments, Nintedanib and Pirfenidone, only show a partial effect by slowing disease progression but do not reverse fibrosis or substantially improve survival [3], underscoring the urgent need for novel therapeutic discovery strategies [6]. One major hurdle for drug discovery is that the bleomycin mouse model fails to recapitulate the progressive nature of the disease and lacks the classic usual interstitial pneumonia (UIP) histopathology [7]. Thus, mouse models have not been very effective at predicting the clinical success of new therapeutics for IPF. The lack of relevant models has also led to a poor understanding of cellular interactions and key signaling pathways involved in the initiation and progression of the disease, which cannot be identified using end-stage patient lung transplant samples available for research. Obtaining samples from patients with interstitial lung abnormalities (ILA) is challenging due to their rarity, sample heterogeneity and the difficulty in predicting disease progression. Precision cut lung slices (PCLS) hold promise for studying the underlying biology of IPF and for developing therapies, but still need improvements in reproducibility, cell survival, and throughput [8].

Given the lack of relevant, reproducible *in vivo* and *ex vivo* models, there has been a push to develop novel *in vitro* models that recapitulate the complex, multicellular, and spatial microarchitecture of progressive fibrosis of the human lung. Traditional monolayer fibroblast cultures, which are still widely used in pharmaceutical research and drug screening pipelines, fail to capture the dynamic crosstalk between alveolar epithelial and mesenchymal cells in three dimensions (3D) [9]. These models do not accurately reflect the cellular and mechanical cues that modulate the initiation and progression of complex diseases, leading to a poor predictive value for determining drug efficacy in patients. While commercial 3D scaffolds like Matrigel provide structural support, they lack alveolar microarchitecture, relevant mechanotransduction biological cues, and the alveolar microenvironment necessary for accurate tissue mimicry. For a complex disease like IPF, a physiologically relevant model requires the creation of the cellular niche with key cell types, such as alveolar type 2 (AT2) epithelial cells and lung fibroblasts, to capture the intricate crosstalk that regulates the underlying biological processes of disease initiation and progression. *In vitro* complex organoid models with multiple cell types typically take a “bottom up” or “top down” approach. For the “bottom up” approach, pluri-or multi-potent cells are provided with matrix, growth factors, and other microenvironmental cues to promote differentiation and self-organization along multiple lineages [10]. A “top down” approach involves combining multiple cell types on a scaffold or matrix and bringing cells together to promote interactions [11–13]. Both approaches are useful and are under development in many organ systems [14]. Primary cells, especially cells from diseased tissue, are ideal for this “top down” approach but are challenging to obtain and culture. Cell lines are an easy solution but often fail to recapitulate important aspects of the disease. The generation of induced pluripotent stem cells (iPSCs) from patients with genetic diseases provides a useful tool to create cell types for the “top down” or “bottom up” approaches and, as they are amenable to gene correction, syngeneic control lines for specific mutations can also be generated, which are important for understanding the biology of the specific mutation being examined [15].

Observational cohort studies show that Familial IPF (FPF) accounts for about 10-20% of patients with IPF and is associated with rare variants in two main biological areas: surfactant proteins (SFTP) and telomere-related genes. In addition, common single-nucleotide polymorphisms (SNPs), such as the MUC5B rs35705950 SNP, likely contribute to the risk of developing IPF. The most common risk variant is the I73T *SFTPC* variant, which encodes a missense mutation and results in altered SFTPC trafficking and proteostasis [16]. Previous work from Alysandratos et al., demonstrated that iPSC-derived AT2 (iAT2) cells with the I73T SFTPC mutation recapitulate the hallmarks of the epithelial dysfunction seen in IPF, but this model lacked an epithelial niche and interactions with other cell types, especially fibroblasts [17].

Single-cell RNA sequencing (scRNA-seq) studies of end-stage IPF lungs have identified diverse cell types and subtypes. One of the key changes is the appearance of K5^-^K17^+^ atypical basaloid epithelial cells that are not present in healthy lungs [18–20]. In addition, specific subtypes of fibroblasts have been described in IPF lungs, specifically fibrotic fibroblasts (expressing high levels of Collagen Triple Helix Repeat Containing 1 (CTHRC1) and Collagen 1A1 (Col1A1)) in fibroblastic foci [21], myofibroblasts (expressing Col1A1 and alpha smooth muscle actin (αSMA)), proliferative fibroblasts (expressing Marker of Proliferation Ki-67 (MKI67)), inflammatory fibroblasts (expressing Interleukin 6 (IL6) and C-C Motif Chemokine Ligand 2 (CCL2) [22], and fibroblasts with a cholesterol metabolism signature (expressing CCAAT Enhancer Binding Protein Beta (CEBPB)) [23, 24]. When modeling IPF, it is ideal to recapitulate the multiple cell types of the disease, the extracellular matrix (ECM) components they produce, and the mechanotransduction forces they impart. The ECM is key for sequestering latent Transforming Growth Factor Beta (TGFβ), which is then available to be activated as a major driver of progressive fibrosis [25]. Thus, the matrix and matrix stiffness are key for modeling IPF *in vitro* [26].

To validate our hypothesis that alveolar microarchitecture, matrix stiffness, and epithelial-fibroblast interactions are essential for modeling fibrogenesis, we developed a human co-culture organoid model, which incorporates primary lung fibroblasts and iAT2 cells from different sources. We developed a human co-culture organoid model, which co-cultures iAT2 cells with the SFTPC I73T mutation (iAT2^I73T/tdT^) with fibroblasts from IPF patients (termed IPF-Fib). We also developed a human co-culture organoid model, which co-cultures the syngeneic corrected control iPSC line (iAT2^tdT/WT^) with fibroblasts from patients without lung disease (termed Healthy-Fib). We seeded these combinations of iAT2 cells and fibroblasts onto microbead scaffolds to mimic cells growing in the interstitial spaces between the 3D alveolar air sacs, thereby recapitulating the lung alveolar niche [11]. We found that the co-culture organoid model with IPF-fib and iAT2^I73T/tdT^ cells produced more extracellular matrix (ECM) than the co-culture organoid model with Healthy-fib and iAT2^tdT/WT^. The more fibrotic cell co-culture combination also produced large amounts of secreted factors and recapitulated aspects of the progressive fibrosis in IPF. We also found that atypical basaloid epithelial cells and fibrotic lung fibroblast cell subtypes appeared over time and resembled the cell subtypes found in advanced-stage IPF tissue. The combination of healthy fibroblasts and iAT2^I73T/tdT^ cells enabled us to examine the initial fibrogenic cues arising from this cellular crosstalk. We also used the cell co-culture organoid model with IPF fibroblasts and iAT2^I73T/tdT^ cells to test the efficacy of antifibrotic drugs that are in the clinic or in development for the clinic. Overall, we showed the utility of our stiff, scaffolded, cell co-culture organoid model for studying cell-cell interactions that drive the initiation of fibrosis and promote its progression and demonstrated the value of our model for drug testing.

## Methods

### Model generation

The model generation involved multiple steps that are explained in a-d:

### **a.** Activation of polyacrylamide microbeads

We procured the polyacrylamide gel beads from Bio-Rad (Cat. 1504130) and activated them following the steps below (i-ii):

i. Benzoquinone solution (0.1M) was added to a coupling buffer (pH 8) in 1:4 ratio. Benzoquinone solution is made up of 1.1g 1,4-p-benzoquinone (Sigma-Aldrich, Cat. 1056504) in a 20% dioxane/water solution. Coupling buffer was made by mixing 47 ml dibasic sodium phosphate solution [2.84 g of sodium phosphate dibasic (Sigma Aldrich, Cat. 4276) in 200ml water) and 3ml of Monobasic sodium phosphate solution (2.4g of sodium phosphate monobasic (Sigma Aldrich, Cat. 4269) in 200 ml of water.
ii. The polyacrylamide beads were mixed in the benzoquinone-coupling buffer solution followed by a series of washing: 2 x 10 minutes with 20% dioxane, 2 x 10 minutes with cell culture grade water, 1 x 20 minutes with sodium acetate buffer (Sodium Acetate – 1164mg, Sodium Chloride – 7.6g, acetic acid – 630ul, Water – 130ml, pH 4.0), 1 x 20 minutes with sodium bicarbonate buffer (Sodium Acetate – 1.68g, Sodium Chloride – 11.6g, Water – 400ml, pH 8.5), and 1x 20 minutes with coupling buffer (pH 7.5). The activated beads were then stored in fresh coupling buffer (pH 7.5) at 4 °C until functionalized.

### **b.** Functionalization of microbeads

A detailed method of functionalization is described in [28]. Briefly, 2.5 ml of the activated beads were rinsed and allowed to soak in 1ml of high-concentration (9.37 mg/ml) rat tail collagen I solution (Corning, Corning, NY, https://www.corning.com) for 6 days at 4 °C. After soaking, the beads were pipetted into a 15-ml centrifuge tube, and the excess collagen I was aspirated, and 8ml of 2mg/ml dopamine hydrochloride (Sigma-Aldrich, H8502) in 50mM Tris buffer (pH 8.5) was added. The tube was rotated at 16.5 rpm on a laboratory rotisserie (Thermo Fisher Scientific Life Sciences, Waltham, MA, https://www.thermofisher.com) for 1 hour at room temperature. Beads were then rinsed 5x in Tris buffer to remove excess collagen, then soaked in experimentally relevant, serum-free media.

### **c.** iAT2 and Fibroblast cell culture and media preparation

Human iAT2 cells were generously provided by Drs. Kotton and Alysandratos labs from Boston University. A detailed method of generating the cells and tdTomato reporter knock-in is described in Alysandratos et al. [29]. Briefly, iAT2s were generated through iPSC-directed differentiation via definitive endoderm into NKX2-1 lung progenitors using previously published methods [29, 48–51]. On days 15-17 of differentiation, live cells were sorted on a high-speed cell sorter (MoFlo Astrios EQ) to isolate NKX2-1+ lung progenitors based on CD47hi/CD26neg gating [52]. Sorted lung progenitors were resuspended in undiluted growth factor-reduced 3D Matrigel (Corning) at a density of 400 cells/µl and distal/alveolar differentiation of cells was performed in CK+DCI medium, consisting of complete serum-free differentiation medium (cSFDM) base supplemented with 3 µM CHIR99021, 10 ng/ml rhKGF (CK), 50 nM dexamethasone (Sigma), 0.1 mM 8-Bromoadenosine 3’,5’-cyclic monophosphate sodium salt (Sigma), 0.1 mM 3-Isobutyl-1-methylxanthine (IBMX; Sigma) (DCI). The resulting epithelial spheres were passaged without further sorting on approximately day 30 (day 28-32) of differentiation, and a brief period (4-5 days) of CHIR99021 withdrawal followed by one week of CHIR99021 addback was performed to achieve iAT2 maturation, as previously shown [48, 50]. After this 14-day period, SFTPCtdTomato+ cells were purified by FACS and resuspended in undiluted growth factor-reduced 3D Matrigel (Corning) at a density of approximately 400 cells/µl with refeeding every other day with CK+DCI medium, according to our published protocol [50]. iAT2 culture quality and purity were monitored at each passage by flow cytometry, with more than 95% of cells expressing SFTPCtdTomato over time, as we have previously detailed[29, 51]. After receiving the cells at UCLA, we reseeded them on Matrigel droplets in the CK-DCI media [53] and cell passages <3 were used for this study.

Primary healthy lung fibroblasts were isolated from distal lung tissue from de-identified healthy donors (D1:65, male, Caucasian, non-smoker; D2:66, F, Caucasian, non-smoker; D3:45, M, Caucasian, Non-smoker) procured from the International Institute for the Advancement of Medicine (IIAM). Primary IPF lung fibroblasts were isolated from transplanted lung tissues from de-identified IPF patients (IPF1: 65, male, Hispanic, previous smoker; IPF2: 70, male, Caucasian, smoker; IPF3: 65, M, Hispanic, smoker) at Ronald Reagan Medical Center, UCLA. Human lung tissue was procured under the UCLA-approved IRB protocol #16-000742. For both healthy and IPF lung tissue, the fibroblasts were isolated in the following way:

The distal tissue was cut into 1cm×1cm pieces and kept in gelatin-coated 6-well plates, submerged in the Fibroblast media (DMEM+F12, 10% FBS, 1% non-essential amino acids, 1% glutamax) for 3-4 weeks to allow the fibroblasts to “crawl out” of the tissue and form an adherent monolayer in the well. The “crawled-out” populations were dissociated with TrypLE (Thermofisher, Cat. 12605036), cultured in cell culture flasks, and passages <5 were used for this study. All fibroblast cultures were exposed to serum-free media (DMEM+F12, 1% non-essential amino acids, 1% glutamax) for 48h before using them for 3D model formation.

### **d.** Bioreactor setting, 3D model formation, and loading into 96-well plates

To develop the 3D model, we used a high aspect ratio vessel (HARV) bioreactor vessel (model: RCCS-4H; Synthecon, Houston, Texas) of 2 ml volume and added 0.5 ml of functionalized microbeads and 1.5 ml of media containing a total of 1 million cells. For an iAT2 and fibroblast (co-culture) model, a mixture of equal parts of serum-free fibroblast and iAT2 media was used. For a fibroblast-only organoid model, only serum-free fibroblast media was used. The vessel was screwed into the bioreactor base, and the beads and the cells were allowed to settle. After sedimentation, the bioreactor was powered on to 4 rpm. For all cellular combinations, after 48h, the cell-coated bead solution was aliquoted 100µl per well in a glass-bottom 96-well plate (Cellvis P96-1.5H-N) with the help of a multichannel pipettor. The 96-well plate was then briefly centrifuged (1000g, 2 min) to settle the cells/ beads at the bottom of the plate and an additional 150µl media was added to each well. The plate was then kept inside an incubator (37°C, 5% CO2, 95%RH) and monitored for the formation of self-organized 3D structures. Within the next 72 hours, the fibroblasts in the model pulled the beads together, and the fully formed 3D models with micro-alveolar structures were observed in each well. From one bioreactor of 2ml capacity, 20 organoids were formed. The number and capacity of the bioreactors were varied as required. For a full 96-well plate, 5×2ml bioreactors or 1×10ml bioreactor were used.

### **e.** Atomic force microscopic analysis

For stiffness measurements, 0, and 6-day old live organoids were transferred to 35mm fluorodishes (WPI, FD35-100) containing phosphate-buffered saline (PBS) buffer and atomic force microscopy was performed at 37 ^0^C in PeakForce Tapping mode using JPK Nanowizerd 4A (Bruker Nano Surface, CA, USA). We used a PeakForce Quantitative Nanomechanics-Live Cell (PFQNM-LC) probe (Bruker AFM probes, CA, USA) with a silicon tip [length=54µm; radius=4.5µm; frequency=45kHz; Spring constant=0.1N/m], specially optimized for soft biological samples. During measurements, multiple (n ≥3) organoid locations were selected and an area of 100µm×100µm was scanned in each location. Force-distance curves were recorded to obtain tumor stiffness. Data analysis was done in JPKSPM Data processing software (version 6)(Bruker, USA). For calculating Young’s modulus, force-distance curves were converted to force-separation curves, and the Hertz-Sneddon model was chosen during model fitting.

### **f.** Immunofluorescence staining, imaging, and image analysis

For surface area measurement, the live organoids were imaged in a Zeiss Inverted Phase Contrast Fluorescence Microscope (Axiovert 40 CFL) in culture days 0, 3, and 6. For image analysis, images of the same set of wells were captured. Each well was divided into four quadrants, and for each quadrant, images were captured at the exact location at each time point in triplicate.

We used ImageJ (National Institute of Health, USA) software to analyze tumor area. For a particular image, the GFP zones were selected and the inbuild area measurement tool of the software was used to calculate area [54]The scale was converted from pixels to microns using the image’s actual scale.

For whole-mount staining, the organoids were first fixed using 4% paraformaldehyde (Thermo Fisher) for 30 minutes at room temperature and then permeabilized using 0.1% TritonX-100 (Sigma-Aldrich) in Tris-buffered saline (TBS) for 15 minutes. After blocking in DAKO (Agilent) for 1h, organoids were incubated with primary antibodies overnight at 4°C. The next day, after triple washing, organoids were incubated in secondary antibodies (ThermoFisher) along with 4’,6-diamidino-2-phenylindole (DAPI) for 1h at room temperature. The following primary antibodies, COL1A1 (Cell Signaling Technology, Cat. 72026, 66948), EpCAM (ab71916), and secondary antibodies Alexa Fluor 488, Alexa Fluor 594, were used. Confocal imaging was performed using Zeiss LSM 880.

For COLA1A fluorescence intensity analysis, we adopted the corrected total cell fluorescence method by ImageJ [55] (For the bulk image analysis for cell count and surface area calculation, we first determined a threshold to separate the background from the foreground regions of each image. We applied Otsu’s method [56], a widely used clustering-based thresholding technique that assumes a bimodal intensity distribution and automatically identifies an optimal threshold value, /_th_.

Using this threshold, we generated a binary mask with the same dimensions as the input image, where pixels with intensities greater than /_th_were labeled as foreground. Positive intensity was then calculated as the mean intensity within the masked (foreground) region. Likewise, negative intensity was computed from the complementary (background) region.

To streamline the process and enable automated analysis across large image batches, we developed a Python-based script that performs these calculations. This automated workflow helps reduce variability and potential human error associated with manual, image-by-image intensity assessment. All analyses were performed using Python version 3.7. The images were stored in grayscale.tif format at a resolution of 1269 × 972 pixels. Examples of intensity and thresholding analysis are shown in Supplemental Figure 1i-j.

### **g.** Luminex assay for secretory biomarker

For Luminex assay, cell culture supernatants were collected, and aliquots were stored at-80 °C. Supernatants were shipped to Eve Technologies (Calgary, Alberta, Canada) on dry ice, and levels of cytokines and chemokines, MMP/TIMPs, and TGFβ1-3s were measured using the Human Cytokine/Chemokine Panel A 48-Plex Discovery Assay (HD48A), Human MMP and TIMP 12-Plex Discovery Assay (HMMP/TIMP-C,O), and TGFB 3-Plex Discovery Assay (TGFβ1-3), respectively.

### **h.** Drug treatment and IC50 calculations

IPF-Fibs (isolated from two IPF patients, IPF1 and IPF2, with demographics as mentioned earlier) with iAT2^I73T/tdT^ cell co-culture organoid models were used for the drug testing experiment. The drugs Nintedanib (Medchem Express, cat. BIBF 1120) and SB-431542 (Medchem Express, cat. HY-10431) were received as powders and diluted in DMSO to a stock concentration of 10 mM. The stock solution was then diluted serially to achieve the final concentrations: 0 (no drug added DMSO control), 1, 10, 30, 50, 80, 100, and 500 nanomolar (nM) for Nintedanib, and 0 (no drug added DMSO control), 1, 10, 30, 50, 100, 500, 1000 nM for SB-431542. The organoids were kept in drug-added media for 6 days with every 48h media change. Brightfield imaging, cell supernatant collection, and Immunofluorescence staining for COLA1 were performed at days 0, 3, and 6. To calculate the IC50, the fluorescence intensity of COLA1 and the percent change in surface area were used as readouts. From the drug concentration and corresponding COLA1 and surface area values, IC50 was calculated using Graphpad Prism (version 10.5) software. All treatments were done in triplicate.

### **i.** Single-cell RNA sequencing fixed RNA profiling

Organoids from four co-culture samples: Healthy-Fib+ iAT2^tdT/WT^, Healthy-Fib+ iAT2^I73T/tdT^, IPF-fib+ iAT2^tdT/WT^, IPF-fib + iAT2^I73T/tdT^, and two fibroblast-only culture samples: Healthy-Fib and IPF-Fib were developed for this experiment. Healthy-Fib was isolated from a healthy donor (65, male, Caucasian, non-smoker), and IPF-Fib was isolated from an IPF patient (IPF1: 65, male, Hispanic, previous smoker). For each model, single cells were collected on day 3 and day 6. triturated 10 times in TrypLE with a 1 mL pipette, followed by a 25-minute incubation at room temperature with trituration every 5 minutes. Afterward, samples were diluted in adDMEM/F12++ and centrifuged at 300x*g* for 5 minutes at 4. The resultant single cell suspensions were then fixed in 4% PFA and stored at-80 prior to preparation of sequencing libraries using the Chromium Next GEM Single-cell Fixed RNA Sample Preparation Kit according to manufacturer instructions (https://www.10xgenomics.com/support/single-cell-gene-expression-flex/documentation/steps/sample-prep/fixation-of-cells-and-nuclei-for-chromium-single-cell-gene-expression-flex). Samples collected after differentiation phase were handled identically with the following exceptions: samples were incubated in a 37 water bath for the TrypLE dissociation and filtered through a 40 µm cell strainer after dilution in adDMEM/F12++. Fixed frozen samples were delivered to UCLA Technology Center for Genomics and Bioinformatics (TCGB) for sequencing library construction following the Chromium Fixed RNA Profiling Reagent Kits user guide for multiplexed samples (https://www.10xgenomics.com/support/single-cell-gene-expression-flex/documentation/steps/library-prep/chromium-single-cell-gene-expression-flex-reagent-kits-for-multiplexed-samples). Libraries were sequenced as 50 base-pair paired-end reads on a NovaSeq X Plus instrument (Illumina).

### **j.** Single-cell RNA sequencing analysis

FRP data were mapped with CellRanger 8.0.0 to the GRCh38 human reference genome. Data was processed and displayed in R using the Seurat, ggplot2, and pheatmap packages. Cells from obviously mistaken library hashing (iAT2 cells in Fib only mixtures) were discarded. Clusters of DEGs were defined using the FindAllMarkers() function, and ontology enrichments were determined using Metascape. Gene networks were determined using geneChorus (https://github.com/Teneth/geneChorus). Gene correlations are calculated in squared scaled space at multiple quality thresholds to determine stable patterns. These patterns are reported as classification of all genes detected into approximately 150 groups for visualization of expression, enrichment, and gene ontology analysis. Gene network enrichment was calculated per cluster and ranked to determine specific and shared characteristics of clusters. Cell-cell communication inference was determined using the iTalk software and database. Pairwise differential gene expression analysis for mixture comparison was calculated with a negative binomial distribution only for Fibroblasts in each sample. Hypoxia Hallmark signal was obtained GSEA-MSigDB database (M5891).

For head to head comparisons of Sample Cluster distributions, the Log ratio was calculated by dividing the frequency in each cluster of sample (A) with the frequency of sample (B), after addition of a pseudocount based on 0.5% of the total cells in both sets, to reduce the effect of small cell groups, and then the Log 2 was taken of the ratio. To show the intensity of comparison differences scaled by the amount of cells involved, the Power Ratio was calculated by multiplying the Log Ratio by the sum of the frequencies plus 10% of the total frequency, divided by the total frequence. The Gene Ontology and biological pathway analysis was done by Metascape [57]. Gene-gene correlation patterns were demarcated using geneChorus v12.3. Gene correlations are calculated in squared scaled space over multiple rounds to determine stable patterns. These patterns are reported as classification of all genes detected into approximately 200 groups for visualization of expression, enrichment, and gene ontology analysis. Cell-cell communication was analyzed by iTALK package in R [58]. For gene class annotation, we cross referenced the Universal Gene Symbol with the AnimalTF database the Human Protein Atlas prediction database.

### **k.** Statistical analysis

All data were compiled from three or more independent replicates for each experimental condition except for the single cell RNA sequencing experiments. Depending on the experiment, data comparisons and statistical significance analyses were performed using Student’s t-test or one-or two-way ANOVA in GraphPad Prism (version 10.5). Significance was defined as p < 0.05. Specific statistical details of the experiments are provided in the Figure legends.

## Materials availability

This study did not generate new unique reagents.

## Results

### The cell co-culture organoid model with IPF fibroblasts and iAT2^I73T/tdT^ cells exhibits matrix deposition, tissue contraction, and loss of iAT2 cells, key hallmarks of progressive fibrosis

In our earlier work, we demonstrated that a polyacrylamide-based stiff hydrogel (elastic modulus: 13kPa) can generate the mechanical cues necessary to model progressive fibrosis in a 2D model [27]. We also previously showed that microbeads can be used as a scaffold to recapitulate the interstitial spaces of the distal lung microarchitecture [11, 28]. In the present work, we modified the 2D 13kPa polyacrylamide scaffold to generate 13kPa polyacrylamide microbeads with an average diameter of 100µm, resembling the diameter of alveolar air sacs, to better recapitulate the stiffness of the alveolar microarchitecture in IPF (Figure 1ai-ii).

The cell co-culture organoid model involves combining 13kPa microbead scaffolds coated with rat tail collagen 1 in a rotating bioreactor with different combinations of fibroblast and iAT2 cell populations for 48 hours. The iAT2 cells were generated from a patient who was heterozygous for the I73T *SFTPC* variant (*SFTPC*^I73T/WT^). A tdTomato fluorescent reporter was knocked into the wild type allele of the *SFTPC* locus and as the tdTomato cassette is followed by a stop/polyA cassette, it prevented expression of the subsequent *SFTPC* coding sequence from the targeted allele, resulting in a *SFTPC*^I73T/tdT^ line [29]. A gene-corrected, syngeneic iAT2 cell line was also generated expressing one allele of the endogenous *SFTPC* locus and the tdTomato reporter (iAT2^tdT/WT^) [29]. For the healthy lung organoid version of the model, we combined healthy age-matched donor-derived fibroblasts at low passage from patients with no prior history of lung disease (Healthy-fib) with the healthy iAT2^tdT/WT^ cells. To develop the cell co-culture organoid with IPF fibroblasts and iAT2^I73T/tdT^ cells, we combined primary, low-passage (<5) IPF patient-derived fibroblasts (IPF-fib) with the iAT2^I73T/tdT^ cells. Figure 1ai shows a schematic of the Healthy-Healthy and IPF Disease-Disease cell combinations in the cell co-culture organoid model. Figure 1aii illustrates functionalized microbeads without cells, the microbeads coated with live cells (Calcein AM intravital dye) showing fibroblasts pulling the cell-coated beads together, and the typical morphology of the fibroblasts (expressing Vimentin) and the iAT2 cells (expressing EpCAM) together on the beads in the cell co-culture organoid model.

The cell types were combined in the bioreactor in a 1:1 cell ratio of iAT2 cells to fibroblasts. We tested different total cell numbers per organoid and then standardized each organoid to a starting density of 100,000 total cells (Supplemental Figure 1a). After 48 hours in the rotary bioreactor, we aliquoted the cell-coated beads into 96-well plates and centrifuged them at low speed to allow the fibroblasts to pull the cell-coated beads together, thereby creating interstitial spaces in which the cells interact. iAT2s were initially cultured in serum-free CK-DCI media containing Keratinocyte Growth Factor (KGF), Dexamethasone, cyclic AMP (cAMP), and the phosphodiesterase inhibitor, 3-isobutyl-1-methylxanthine (IBMX). The fibroblasts were initially cultured in Dulbecco’s Modified Eagle Medium (DMEM) supplemented with 10% fetal bovine serum (FBS). For the co-culture organoid model, we combined these two media in a 1:1 ratio and removed the FBS from the fibroblast media. Therefore, the media used for all experiments was CK-DCI media combined with serum-free DMEM in a 1:1 ratio.

IPF is characterized by aberrant deposition of ECM proteins, and Collagen 1 is one of the major components of this ECM [30, 31]. Loss of epithelial cells is also a hallmark of remodeling in IPF tissue. We therefore performed immunofluorescence (IF) staining for Collagen 1A1 (COL1A1) and Epithelial Cell Adhesion Molecule (EpCAM) to identify epithelial cells in the cell co-culture organoid models on day 6 of culture when remodeling is occurring (Figures 1b,c). The cell co-culture organoid model with the Healthy-Fib and healthy iAT2^tdT/WT^ cells demonstrated a uniform distribution of COL1A1 across the scaffold with epithelial cells (Figure 1b), while the cell co-culture organoid model with the IPF-Fib and iAT2^I73T/tdT^ showed larger areas of COL1A1 IF staining, and loss of EpCAM^+^ cells (Figure 1c). Quantification of the fluorescence intensity of COL1A1 IF staining per cell showed more collagen in the IPF-Fib and iAT2^I73T/tdT^ cell co-cultures when compared with the Healthy-Fib and iAT2^tdT/WT^cell co-culture (p<0.001) (Figure 1d), and less EpCAM^+^ cells (p<0.001)(Figure 1e). We also imaged the SFTPC tdTomato reporter to assess the number of SFTPC-expressing cells in the two models and we found that the IPF-Fib and iAT2^I73T/tdT^ cell co-culture organoid model exhibited significantly fewer tdTomato^+^ cells than the Healthy-Fib with the healthy iAT2^tdT/WT^cell co-culture organoid model (Figures 1f, g).

**Figure 1.**
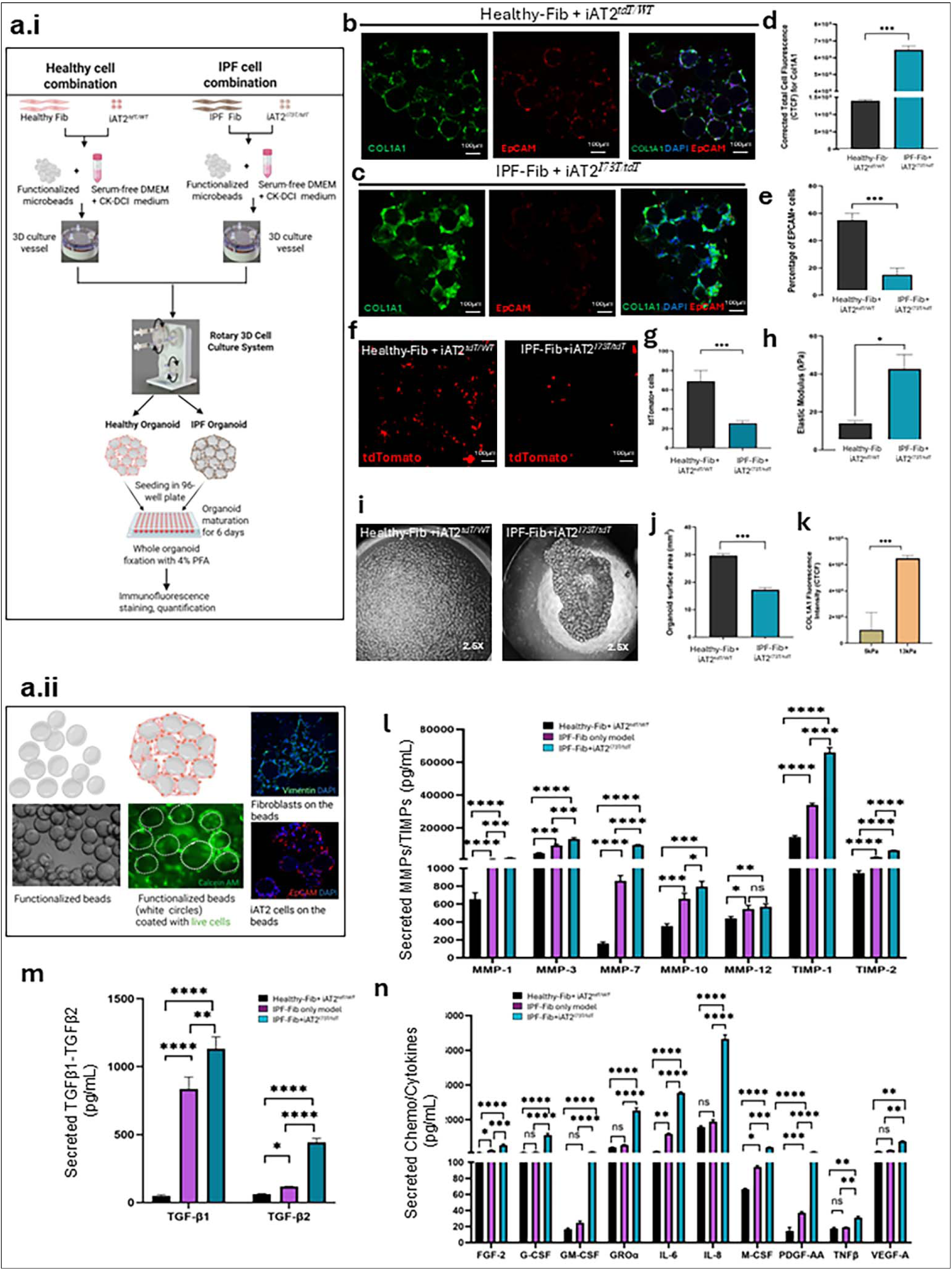
Development of Healthy-Fib with iAT2^tdT/WT^ cell and IPF-Fib with iAT2^I73T/tdT^ cell co-culture organoid models. a.i. Schematic of steps for model generation, timeline, and readouts. Healthy cell combination consists of primary human donor lung fibroblast and syngeneic corrected iPSC-derived AT2 cells (iAT2^tdT/WT^); IPF cell combination consists of primary human IPF patient lung fibroblast and iPSC-derived AT2 with SFTPC 173T mutation (iAT2^I73T/tdT^). **a.ii.** Graphics and corresponding images of functionalized microbeads before cell-coating, live-cell-coated (labeled with Calcein AM viability stain) aggregated microbeads pulled together by the fibroblasts, and morphology of fibroblasts (Vimentin), and iAT2 cells (EpCAM) in the cell co-culture organoid models. **b.** Immunofluorescent staining on Day 6 for COL1A1(green) and EpCAM (red) expression in the Healthy-Fib with iAT2^tdT/WT^ cell co-culture organoid models (biological replicate n=2). Scale bar = 100µm. **c.** Day 6 COL1A1 and EpCAM expression in the IPF-Fib with iAT2^I73T/tdT^ cell co-culture organoid model (biological replicates n=2). Scale bar = 100µm. **d.** Day 6 COL1A1 CTCF (corrected total cell fluorescence) [54] intensity quantification in the Healthy-Fib with iAT2^tdT/WT^ cell and IPF-Fib with iAT2^I73T/tdT^ cell co-culture organoid models. P values are calculated from technical replicates (n=3) for each biological replicate using one-way ANOVA ***: P<0.001 **e.** Percentage of EpCAM^+^ cells in Healthy-Fib with iAT2^tdT/WT^ cell and IPF-Fib with iAT2^I73T/tdT^ cell co-culture organoid models on day 6. P values are calculated from technical replicates (n=3) for each biological replicate using one way ANOVA ***: P<0.001. **f.** tdTomato+ iAT2 cells in Healthy-Fib with iAT2^tdT/WT^ cell and IPF-Fib with iAT2^I73T/tdT^ cell co-culture organoid models on day 6. Scale bar = 100µm. **g.** Quantification of tdTomato+ cells. P values are calculated from technical replicates (n=3) using Student’s t-test ***: P<0.001. **h.** Quantification of Elastic Modulus of Healthy-Fib with iAT2^tdT/WT^ cell and IPF-Fib with iAT2^I73T/tdT^ cell co-culture organoid models on day 6. P values are calculated from technical replicates (n=3) for each biological replicate (n=2) using Student’s t-test. *: P<0.05. **i.** Brightfield images of Healthy-Fib with iAT2^tdT/WT^ cell and IPF-Fib with iAT2^I73T/tdT^ cell co-culture organoid models on day 6. **j.** Surface area quantification on day 6, P values are calculated from technical replicates (n=3) for each biological replicate (n=3) using Student’s t-test ***: P<0.001. **k.** COL1A1 CTCF quantification of the IPF-Fib with iAT2^I73T/tdT^ cell co-culture organoid model seeded on 5kPa and 13kPa scaffolds on day 6, P values are calculated from technical replicates (n=3) for each biological replicate (n=2) using Student’s t-test, ***: P<0.001. **l-n:** Luminex assay of panels of secreted markers in the Healthy-Fib with iAT2^tdT/WT^ cell co-culture organoid model, IPF-Fib with iAT2^I73T/tdT^ cell co-culture organoid model and IPF-Fib only organoid culture models on day 6 from conditioned media, **l.** MMPs and TIMPs **m.** TGFβ1 and TGFβ2, and **n.** secreted chemokines/cytokines. P values are calculated from technical replicates (n=3) for each biological replicate (n=1) using one-way ANOVA for each biomarker. ***: P<0.001, **: P<0.01, *: P<0.05.

Mechanical properties of tissues drive cellular behavior, and the stiffness of tissues is largely derived from matrix deposition [32]. Therefore, to assess the stiffness of the organoid models, we next measured whether the IPF-Fib and iAT2^I73T/tdT^ cell co-culture organoid model and the Healthy-Fib with the iAT2^tdT/WT^ cell co-culture organoid model altered the stiffness of the scaffolds (13kPa) they were seeded on. Atomic Force Microscopy measurements revealed that the matrix stiffness increased substantially for the IPF-Fib and iAT2^I73T/tdT^ cell co-culture organoid model (42.66kPa) within the 6-day culture period. In contrast, the stiffness remained almost unchanged for the Healthy-Fib with the healthy iAT2^tdT/WT^ cell co-culture organoid model (13.96kPa)(Figure 1h).

Increased production of ECM and increased contractility of fibroblasts are responsible for scarring in IPF, resulting in a reduction in lung volume [33]. We therefore examined contractility in the cell co-culture organoid models by measuring the surface area of each organoid in each well. In the Healthy-Fib with the healthy iAT2^tdT/WT^ cell co-culture organoid model, we calculated an average of 9% decrease in surface area due to the contractility of fibroblasts over the 6 days in culture, while in the IPF fibroblasts and iAT2^I73T/tdT^ cell co-culture organoid model, we noticed an average of 46% decrease in average surface area over the same 6-day period (Figures 1i, j). Taken together, we have optimized the IPF-Fib and iAT2^I73T/tdT^ cell co-culture organoid model and the Healthy-Fib and healthy iAT2^tdT/WT^ cell co-culture organoid model and shown that the IPF-Fib and iAT2^I73T/tdT^ cell co-culture organoid model expressed more COL1A1, showed more remodeling with a reduced surface area, and the presence of fewer iAT2 cells.

### Scaffold stiffness drives progressive fibrosis in the cell co-culture organoid model

As the cell co-culture organoid model on stiff 13kPa microbeads induced collagen deposition and loss of iAT2 cells (Figure 1b-h), we next assessed collagen production and iAT2 cell retention in a system with low stiffness. Here, we used Matrigel instead of 13kPA microbeads as a non-stiff scaffold comparison to assess cell-intrinsic fibrotic responses. We combined iAT2^I73T/tdT^ with IPF fibroblasts (IPF cell co-culture combination) and iAT2^tdT/WT^ with healthy fibroblasts (healthy cell co-culture combination) in Matrigel, by mixing iAT2 cells and Fibroblasts in a 1:1 ratio and keeping a total of 10,000 cells/Matrigel droplet. We then analyzed COL1A1 expression and tdTomato+ iAT2 cell numbers (Supplemental Figures 1b-e). We observed that the IPF cell co-culture combination induced higher COL1A1 expression and fewer tdTomato+ iAT2 cells compared to the healthy cell combination in Matrigel droplets. This suggests that IPF fibroblasts and/or iAT2^I73T/tdT^ cells have an intrinsic ability to generate more COL1A1 irrespective of matrix stiffness. However, this fibrotic phenotype was not as robust as that seen in the 13kPa 3D microbead scaffold models, where the same IPF cell co-culture combinations demonstrated significantly higher (p<0.0001) COL1A1 expression than the IPF cell co-culture combinations in Matrigel (Supplemental Figure 1f). We therefore surmised that the scaffold’s stiffness drives the fibrotic phenotype of the cell co-cultures.

To test this further, we used low-stiffness (5kPa) or high-stiffness (13kPa) microbeads in cell co-cultures. We seeded IPF fibroblasts with iAT2^I73T/tdT^ on 5kPa or 13kPa microbead scaffolds and found that the COL1A1 expression was 6-fold higher in the 13kPa scaffold IPF co-culture organoid model as compared to the 5kPa model (Figure 1k, Supplemental Figure 1g). Overall, these data demonstrate that scaffold stiffness plays a critical role in collagen production and iAT2 cell number in our cell co-culture organoid models. Based on this data, we moved forward with cell co-culture organoid models with a 13kPa microbead scaffold.

### Fibroblast-only and fibroblast + iAT2 cell co-culture organoid models have different secretory profiles

Given the increased collagen production and reduced iAT2 cell numbers in the IPF-Fib with iAT2^I73T/tdT^ cell co-culture organoid model, we next examined differences in secreted factors from the iAT2^I73T/tdT^ cells with the IPF fibroblasts combination model, as compared to the IPF fibroblast-only model. Factors associated with disease progression in IPF include Matrix Metalloproteinases (MMPs) and their inhibitors (TIMPs), chemokines, cytokines, and TGF-β1 and TGF-β2. We therefore quantified these proteins from conditioned media to compare the models. We cultured IPF fibroblasts only (IPF-Fib-only organoid model), co-cultures of IPF-Fib with iAT2^I73T/tdT^ cells, and co-cultures of Healthy-Fib with iAT2^tdT/WT^ cells on the stiff (13kPa) polyacrylamide microbead scaffold and then performed a Luminex secretory panel for these factors on the supernatant from the day 6 cultures (Figure 1l-n). Interestingly, except for MMP-12, all the biomarkers were significantly higher in the organoid co-cultures of IPF-Fib with iAT2^I73T/tdT^ cells than in the IPF-Fib-only organoid cultures, and organoid co-cultures of Healthy-Fib with iAT2^tdT/WT^ cells. Moreover, several chemokines and cytokines (G-CSF, GM-CSF, GRO-α, IL-8, TNF-β, and VEGF-A) showed no significant difference between the organoid co-cultures of Healthy-Fib with iAT2^tdT/WT^ cells and IPF-Fib-only organoid cultures, but were significantly increased in the organoid co-cultures of IPF-Fib with iAT2^I73T/tdT^ cells. This demonstrates that the epithelial-mesenchymal cell interactions in the organoid co-cultures of IPF-Fib with iAT2^I73T/tdT^ cells elicit cellular changes that drive the secretion of factors that could be important for the progression of fibrosis in the model.

### The organoid co-cultures of IPF fibroblasts with iAT2^I73T/tdT^ cells recapitulate the cellular diversity and cell subtypes characteristic of IPF

In addition to the cell co-culture combinations generated in the organoid co-cultures of Healthy-Fib with iAT2^tdT/WT^ cells and organoid co-cultures of IPF-fib with iAT2^I73T/tdT^ cells (Figures 1b,c), we also generated organoids from other cell combinations. We co-cultured healthy fibroblasts with iAT2^I73T/tdT^ mutant cells, and IPF Fibroblasts with iAT2^tdT/WT^ syngeneic corrected cells to examine the phenotypes from these healthy-diseased, epithelial-mesenchymal cell combinations. In the organoid co-cultures of Healthy-Fib with iAT2^tdT/WT^ cells, the beads were loosely packed, while in organoid co-cultures of IPF-fib with iAT2^I73T/tdT^ cells, the beads were closely packed and surrounded by dense matrix. However, the combinations of Healthy Fibroblasts with iAT2^I73T/tdT^ cells and IPF Fibroblasts with iAT2^tdT/WT^ cells developed a fibrotic phenotype that appears to be intermediate between that of the organoid co-cultures of IPF-fib with iAT2^I73T/tdT^ cells and organoid co-cultures of Healthy-Fib with iAT2^tdT/WT^ cells (Figure 2a, Supplemental Figure 2a). To investigate the interactions between the iAT2 cells and fibroblasts in the cell co-culture organoid models that are driving these phenotypes, we performed single-cell RNA sequencing (scRNA-seq) of the organoid co-cultures of Healthy-Fib with iAT2^tdT/WT^ cells, the organoid co-cultures of IPF-Fib with iAT2^I73T/tdT^ cells, as well as from the models of combinations of Healthy-Fib with iAT2^I73T/tdT^ cells, and IPF-Fib with iAT2^tdT/WT^ cells (Figure 2b). We also performed scRNA-seq of two fibroblast-only organoid models, IPF-Fib and Healthy-Fib, to examine how iAT2 cells alter the distinct fibroblast gene expression patterns (Figure 2b). The model requires fibroblasts to maintain the scaffold; thus, cultures of iAT2s only could not be established. The six samples were prepared simultaneously, with all parameters (scaffold material, cell number, and cell passage) held constant to minimize experimental variability. To mimic early-and later-stage disease, cells were collected In the Uniform Manifold Approximation and Projection (UMAP), iAT2s and Fibroblasts were clearly separated from each other (inset in Figure 2c). Cells derived from IPF-Fib only organoids and Healthy-Fib only organoids were separated on the fibroblast region of the UMAP (Figure 2c). The cells from the co-culture organoid models on the UMAP showed some overlap with the cells from the fibroblast-only organoid models, but they also showed the emergence of new clusters. This allowed us to label different cell clusters based on known markers of specific cell subtypes in IPF tissue (Figure 2c,d)[17, 34–38]. The iAT2 cell clusters included the SFTPC^high^ cluster, the SFTPC^low^ cluster, the transitioning Secretoglobin Family 3A Member 2 (SCGB3A2)-expressing cluster, the Epithelial Membrane Protein 2 (EMP2)-expressing cluster, the Keratin (K)5^-^K17^+^atypical basaloid cell cluster, and the Marker of Proliferation Ki-67 (MKI67^+^) proliferating iAT2 cell cluster (Figure 2d).

**Figure 2.**
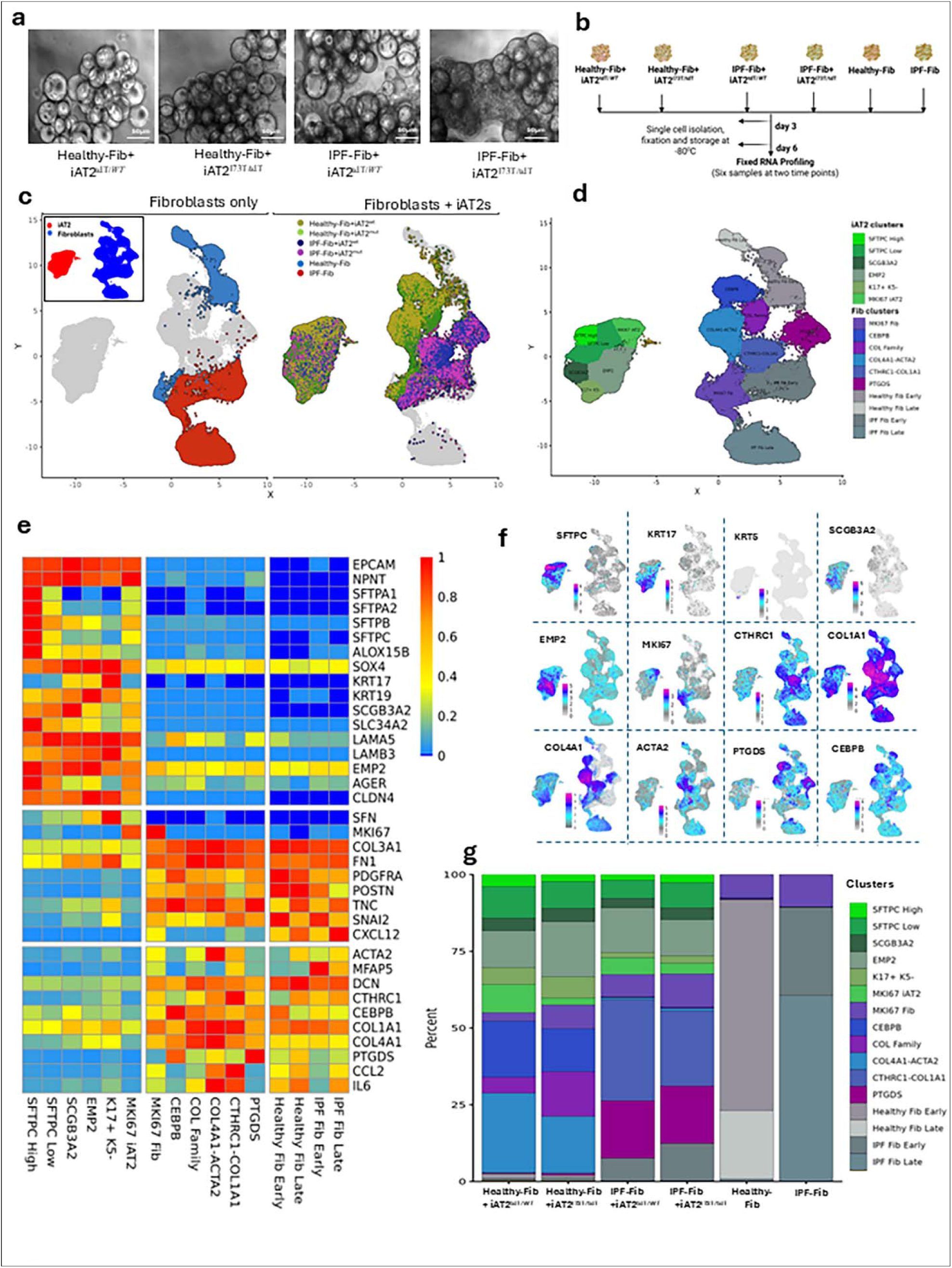
iAT2 cell-Fibroblast interactions are recapitulated in the cell co-culture organoid models. **a.** Changes in the phenotype of the model as seen with brightfield microscopy with different cell combinations on day 6 of culture, Scale bar = 50µm. **b.** Schematic of the different samples, timepoints and models used for scRNA-seq. **c.** UMAP of iAT2 and fibroblast cell co-cultures organoid models and fibroblast-only organoid models. The inset shows iAT2 cells versus fibroblasts transcriptomic group locations in the UMAP. **d.** Assignment of cell clusters based on gene expression. **e.** Heatmap showing the combined gene expression signatures in the fibroblast with iAT2 cell co-culture organoid models and fibroblast-only organoid models. **f.** Percent distribution of cell clusters based on fibroblast with iAT2 cell co-culture organoid models and fibroblast-only organoid models. Clusters are color coordinated with **d**. **g.** UMAPs of specific genes expressed in the cell clusters. at two time points (day 3 and day 6) for scRNA-seq with the Fixed RNA Profiling FLEX assay (10X Genomics).

The fibroblast clusters that were enriched in the fibroblast and iAT2 cell co-culture organoid models were based on fibroblast gene expression patterns seen in IPF tissue. These clusters from the co-culture organoid models included the Prostaglandin D2 Synthase (PTGDS) inflammatory fibroblast cluster, the lipid metabolism CCAAT Enhancer Binding Protein Beta (CEBPB^+^) cluster, the Collagen (COL) family cluster (Col1A1^+^, Col5A1^+^, Col8A1^+^, Col12A1^+^), the α-Smooth Muscle Actin (ACTA2^+^)/Col4A1^+^myofibroblast cluster, and the Collagen Triple Helix Repeat Containing 1 (CTHRC1^+^)/Col1A1^+^ fibrotic cell cluster (Figure 2d) [18–20]. The MKI67^+^ proliferating fibroblast cluster was present in both the fibroblast-only organoids and the fibroblasts with iAT2 cell co-cultured organoids (Figure 2c,d). Different cell types from IPF tissue can be distinguished based on a few specific more highly expressed gene markers, and we therefore examined the fibroblast and iAT2 cell clusters from the cell co-culture organoid models for these predicted gene expression differences across the clusters, and these are represented by a heat map and UMAPs (Figure 2e,f, Supplemental Figure 2b). We found subtle yet distinct gene expression patterns in each epithelial and fibroblast cluster, with the greatest difference being between the fibroblast and epithelial cell clusters themselves. The K5^-^K17^+^atypical basaloid cell cluster was associated with loss of Surfactant expression and an increase in SRY-box transcription factor 4 (SOX4), Claudin 4 (CLDN4), Fibronectin (FN1) and Stratifin (SFN). We also analyzed the top 10 differentially expressed genes (DEGs) of each cell cluster in an unbiased manner from the cell co-culture organoid models and identified that the K5^-^K17^+^ atypical basaloid epithelial cells also express PDGFB, IL1A, WNT7A, and MMP10 (Supplemental Figure 2c), which are all involved in a complex pathway of injury, repair and ECM remodeling. We also found that CTHRC1^+^COL1A1^+^ fibroblasts express Twist2, Robo2, and CXCL3, which are involved in cell migration, development and ECM remodeling.

We also examined enriched gene expression networks and identified specific gene ontologies (GO) for each cell cluster (Supplemental Figure 2d). Both the healthy and IPF fibroblast-only organoid cultures expressed genes related to development and ECM (Supplemental Figure 2d), but the healthy lung fibroblast-only cultures also expressed genes related to the interferon response (Supplemental Figure 2d, Clusters 3,5), and the IPF fibroblast-only cultures also expressed genes related to cell motility (Supplemental Figure 2d, Clusters 6,11). Within the iAT2-derived epithelial cell clusters, the K5^-^K17^+^ atypical basaloid cell cluster showed GO terms related not only to epidermal development but also to Matrisome-related genes and RHO GTPase signaling (Supplemental Figure 2d, Cluster 16). The transitioning cell clusters that expressed SCGB3A2 or EMP2 both expressed cell junction and tight junction-related genes (Supplemental Figure 2d, Clusters 12, 15). The CTHRC1^+^ cell Cluster 8 GO terms are related to Golgi vesicle transport and collagen formation, while the Col4A1^+^/ACTA2^+^ myofibroblast Cluster 10 GO terms show the Matrisome and matrix organization. The COL family cell Cluster 9 GO terms relate to Matrisome, ECM remodeling, and development. In addition to prostaglandin signaling, the PTGDS cell Cluster 7 GO terms include Wnt signaling, while the CEBPB cell Cluster 4 GO terms relate to cholesterol biosynthesis. Therefore, GO terms from IPF fibroblasts co-cultured with iAT2 cells in the organoid model (Supplemental Figure 2d, Clusters 1,2,4,7-10,12-16) show that there are clusters of fibroblasts with gene expression patterns that are reminiscent of fibroblast subtypes seen in IPF, and which are not observed in the GO terms from the IPF fibroblast-only organoid cultures [18, 19, 21, 36].

Notably, we also examined the proportions of each of the cell clusters across the different cell culture organoid models and found them to be different (Figure 2g). The organoid co-cultures of Healthy-Fib with iAT2^tdT/WT^ cells or iAT2^I73T/tdT^ cells predominantly gave rise to the Col4A1^+^/ACTA2^+^ myofibroblast cluster, the COL family cluster, and the CEBPB cluster, while the organoid co-cultures of IPF-Fib with iAT2^I73T/tdT^ cells or iAT2^tdT/WT^ cells predominantly gave rise to the CTHRC1^+^/Col1A1^+^ fibrotic cell cluster and the PTGDS inflammatory fibroblast cluster. Interestingly, the presence of iAT2^I73T/tdT^ cells or iAT2^tdT/WT^ cells in the co-culture organoid models did not change the predominant subtypes of fibroblasts seen in each co-culture organoid model. These cell co-culture organoid model cluster proportions differ markedly from those observed in fibroblast-only organoid models (Figure 2g). This suggests that the IPF fibroblasts intrinsically increase their fibrotic (CTHRC1 cluster) and inflammatory (PTGDS cluster) programs in co-culture with iAT2 cells, while healthy fibroblasts seem to respond to the stiff matrix of the bead scaffold by upregulating collagens (COL Family cluster) and ACTA2 (myofibroblast cluster) in response to co-culture with iAT2 cells (Figure 2g).

Taken together, the cellular diversity in the organoid co-cultures of IPF-Fib with iAT2^I73T/tdT^ cells recapitulates some of the cellular components of IPF tissue, including the K5^-^K17^+^ atypical basaloid cells, CTHRC1^+^Col1A1^+^ fibroblasts, and PTGDS-expressing fibroblasts. This creates a more representative model of IPF than the fibroblast-only organoid cultures. We therefore next sought to examine how the diversity of the cell clusters changes over time in the cell co-culture organoid models.

### The proportions of each cell cluster change over time in the cell co-culture organoid models

To further understand the evolution of gene expression changes between day 3 and day 6 of culture in the cell co-culture organoid models, we analyzed scRNA-seq data to identify clusters of both fibroblast and epithelial cell types at day 6 compared to day 3 of culture (Figure 3a). Specifically, we quantified the proportions of cells in different clusters in each co-culture organoid model at days 3 and 6 (Figure 3a, Supplemental Figure 3a). We found that SFTPC-expressing iAT2 cells are reduced on day 6 compared with day 3, while K5^-^K17^+^ expressing iAT2 cells emerged on day 6. Similarly, the Col4A1^+^ACTA2^+^ and the CTHRC1^+^Co1A1^+^ cell clusters also emerged on day 6 (Figure 3a). On the other hand, the Col family cell cluster was observed in healthy lung fibroblast co-culture organoids on day 3 and was no longer present on day 6 (Figure 3a). The UMAP in Figure 3b further shows the change in cell clusters at day 3 and day 6 of culture. Overall, our cell co-culture organoid models recapitulate many dynamic cell types seen in IPF, and therefore, we next interrogated whether these models can be used to study complex cell-cell interactions that may initiate and drive progressive fibrosis.

**Figure 3.**
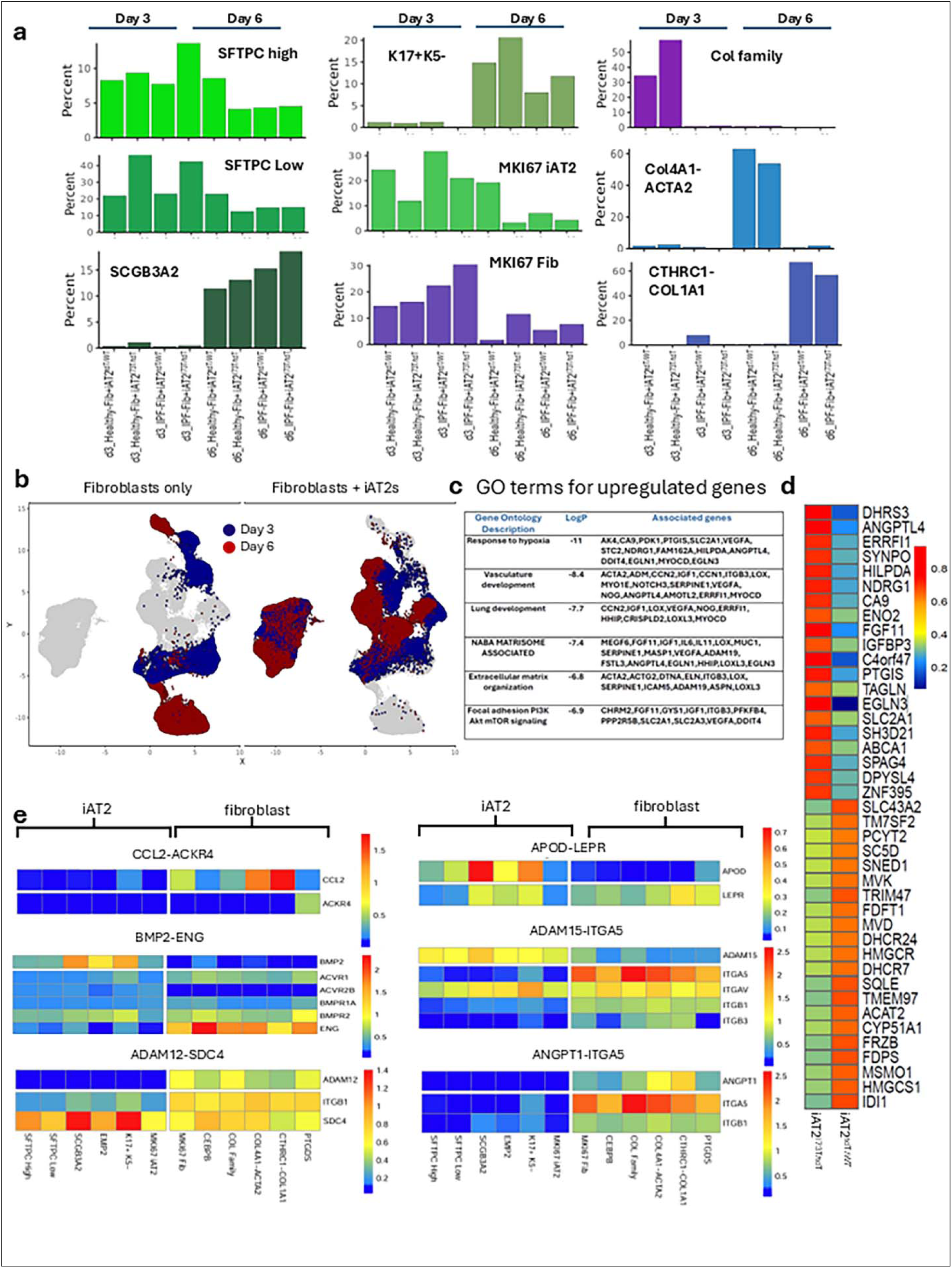
Cellular interactions and disease progression stages are recapitulated in the cell co-culture organoid models: **a.** Percent quantification of clusters per sample on day 3 and day 6. **b.** UMAP with color-coded day 3 (blue) and day 6 (red) gene locations. **c.** Gene Ontology (GO) terms associated with the co-culture combination of Healthy fibroblasts and iAT2^I73T/tdT^ cells. **d.** Displayed are the top 20 highest-and lowest-fold change genes in Healthy fibroblasts when cocultured with iAT2^I73T/tdT^ cells vs. iAT2^tdT/WT^. **e.** Receptor-ligand interactions identified for CCL2-ACKR4, APOD-LEPR, BMP2-ENG, ADAM12-SDC4, ADAM15-ITGA5 and ANGPT1-ITGA5.

### The Healthy-Fib with iAT2^I73T/tdT^ cell co-culture organoid model uncovers cell-cell interactions involved in the initiation and progression of fibrotic remodeling

It is challenging to study the earliest drivers of human lung fibrosis, as it is not possible to obtain patient tissue at the onset of IPF before any clinical phenotype is present. Our model enables the study of the earliest iAT2 cell-fibroblast interactions. To understand how iAT2^I73T/tdT^ cells might induce cellular changes in healthy fibroblasts, we asked how the gene expression state of healthy fibroblasts differs when they are co-cultured with iAT2^tdT/WT^ versus iAT2^I73T/tdT^ cells. We specifically compared differential gene expression in healthy fibroblasts co-cultured with iAT2^I73T/tdT^ cells versus iAT2^tdT/WT^ cells at day 3 to capture early transcriptional responses prior to overt matrix deposition. This comparison showed increased expression of genes in healthy lung fibroblasts co-cultured with iAT2^I73T/tdT^ cells that are involved in the hypoxic response, the Matrisome, and development, including ANGPTL4, DHRS3, EGLN3, ENO2, IL-6, ACTA2, TGLN, and LOXL3 (Figure 3c,d). This suggests that when the healthy fibroblasts encounter iAT2^I73T/tdT^ cells their interaction induces genes involved in matrix production and development as well as hypoxia response genes. In contrast, genes that are induced in healthy fibroblasts when they first interact with iAT2^tdT/WT^ cells are enriched for GO terms related to lipid metabolism, which may reflect their role in supporting surfactant metabolism (Supplemental Figure 3b).

### Single-cell RNA-seq of the cell co-culture organoid models uncovers receptor-ligand interactions between the cell clusters

To better understand the cellular interactions in the cell co-culture organoid models, we performed receptor-ligand analysis of the scRNA-seq data (Figure 3e, Supplemental Figure 3c). We found that the CTHRC1^+^, COL1A1^+^ cells express CCL2, and its ACKR4 receptor is expressed only on the PTGDS fibroblast cluster (Figure 3e). Apolipoprotein D (APOD) is expressed by the SCGB3A2^+^ iAT2 cells and its Leptin Receptor (LEPR) is found on CTHRC1^+^ fibroblasts (Figure 3e). The BMP2 ligand is expressed by SCGB3A2^+^, EMP2^+^ and K5^-^K17^+^expressing iAT2s and its Endoglin (ENG) receptor by all the fibroblast clusters, but especially the CEBPB cluster (Figure 3e). The a disintegrin and metalloproteases (ADAM)15 ligand is predominantly expressed by iAT2 cells, and the fibroblast clusters express its Integrin alpha (ITGA)5 receptor (Figure 3a).

The ligand ADAM12 is only expressed by the fibroblast cell clusters and its receptor Syndecan (SDC)4 is expressed by the epithelial cells, but especially SCGB3A2^+^ and K5^-^K17^+^ expressing iAT2 clusters (Figure 3e). Angiopoietin 1 (ANGPT1) is only expressed by the fibroblast clusters as there are no endothelial cells in the model, and the ITGA5 receptor is more highly expressed on Col4A1^+^-ACTA2^+^ fibroblast and COL family fibroblast clusters (Figure 3e). Most of these ligands and receptors have been described in IPF or fibrosis, indicating that our model is replicating several of the cell-cell interactions that are known in IPF tissue [39–41].

### The IPF-Fib with iAT2^I73T/tdT^ cell co-culture organoid model can be used to evaluate the efficacy of Nintedanib and the Alk5 inhibitor (SB-431542)

To examine whether our IPF-Fib with iAT2^I73T/tdT^ cell co-culture organoid model will respond to the FDA-approved therapy Nintedanib (Medchem Express, cat. BIBF 1120) and the TGFβ inhibitor SB-431542 (Alk5 inhibitor) (Medchem Express, cat. HY-10431), we tested these drugs in the cell co-culture organoid models using lung fibroblasts derived from two IPF patients in combination with the iAT2^I73T/tdT^ cells. We performed an 8-point dose-response (Figure 4a-d, Supplemental Figure 4a,b). We used a non-linear log(inhibitor) vs. response curve method [42] for two endpoint read-outs: COL1A1 fluorescent intensity and the percent reduction of organoid surface area, to calculate the half-maximal inhibitory concentration (IC50)(Figure 4a-d, Supplemental Figure 4a,b). Nintedanib concentrations were: 1, 10, 30, 50, 80, 100, and 500 nanomolar (nM), and SB-431542 concentrations were: 1, 10, 30, 50, 100, 500, and 1000 nM.

**Figure 4.**
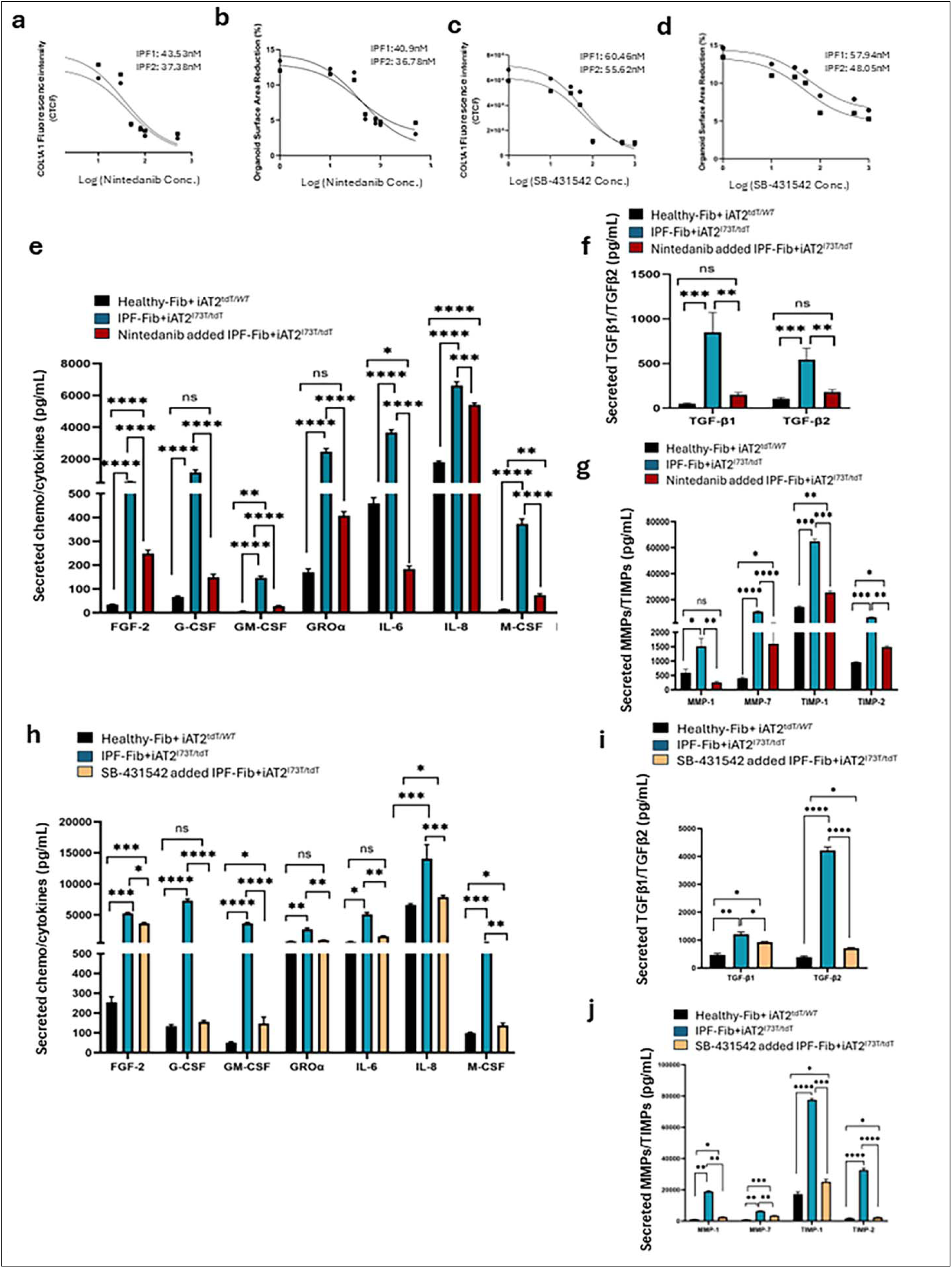
Application of IPF-Fib with iAT2^I73T/tdT^ cell co-culture organoid model for drug testing: a-b: IC50 values of Nintedanib-treated IPF-Fib with iAT2^I73T/tdT^ cell co-culture organoid model developed from IPF-Fib isolated from two different IPF patients (IPF1 and IPF2, see methods for demographics). IC50 was calculated by **(a)** COL1A1 expression changes and **(b)** organoid surface area reduction. **c-d:** IC50 values of SB-431542-treated IPF-Fib with iAT2^I73T/tdT^ cell co-culture organoid model developed from IPF-Fib isolated from two different IPF patients (IPF1 and IPF2) calculated by **(c)** COL1A1 expression changes and **(d)** organoid surface area reduction. **e-g**: Luminex assay of panels of secreted markers in the Nintedanib-treated (40nM, IC50 value for IPF1) and untreated IPF-Fib with iAT2^I73T/tdT^ cell co-culture organoid models, and untreated Healthy-Fib with iAT2^tdT/WT^ cells co-culture organoid models: **e.** MMPs and TIMPs, **f.** TGFβ1 and TGFβ2, and **g.** chemokines and cytokines. **h-j**: Luminex assay of panels of secreted markers in the SB-431542-treated (60nM, IC50 value for IPF1) and untreated IPF-Fib with iAT2^I73T/tdT^ cell co-culture organoid models, and untreated Healthy-Fib with iAT2^tdT/WT^ cells co-culture organoid models **h.** MMPs and TIMPs, **i.** TGFβ1 and TGFβ2, and **j.** chemokines and cytokines. P values are calculated from technical replicates (n=3) for each biological replicate (n=1) using one-way ANOVA for each biomarker. ***: P<0.001, **: P<0.01, *: P<0.05.

The dose-dependent responses for both Nintedanib and SB-431542 with COL1A1 fluorescent intensity (Figures 4a,c) and percent surface area reduction (Figure 4b,d) showed that the IC50 values calculated for the two patients (IPF1 and IPF2) from the two read-outs were in close agreement. The small IC50 difference between the two patients suggests that the model may capture inter-patient variability and may be able to assess individual patient responses.

We also examined the autotaxin inhibitor, Ziritaxestat, which only showed a response at 10nM and not at 1μM or 100nM in the organoid co-cultures of IPF-Fib with iAT2^I73T/tdT^ cells (Supplemental Figure 4c, e). The phosphodiesterase E4B inhibitor showed a response at 1μM and 10nM, but not at 100nM, but the CK-DCI media contains a non-specific phosphodiesterase inhibitor, which may confound these results (Supplemental Figure 4d, e). Notably, none of the compounds tested, even at 1μM, showed a full response comparable to the Healthy-Fib with iAT2^tdT/WT^ cell co-culture organoid model.

To further investigate the effects of Nintedanib and SB-431542, we examined the secreted biomarker panel of the drug-treated cell co-culture organoid model of IPF-Fib with iAT2^I73T/tdT^ cells (derived from patient IPF-1) and compared this with the secreted biomarkers from the non-drug-treated IPF-Fib with iAT2^I73T/tdT^ cell co-culture organoid model, and the non-drug-treated cell co-culture organoid model of Healthy-Fib with iAT2^tdT/WT^ cells. The effect of nintedanib on TGFβ1, TGFβ 2, acute inflammatory factors, and MMPs/TIMPs is shown in Figure 4e-g, and the effect of SB-431542 is demonstrated in Figure 4h-j. Among the chemokines and cytokines, both drugs lowered the secreted protein levels of G-CSF and GRO-α of the cell co-culture organoid model of IPF-Fib with iAT2^I73T/tdT^ cells to a level similar to that seen in the cell co-culture organoid model of Healthy-Fib with iAT2^tdT/WT^ cells, while neither of the drugs showed a significant effect on FGF-2 (Figure 4e, h). Interestingly, both the inflammatory factors IL-6 and IL-8 were reduced by SB-431542, whereas they were not reduced as much by Nintedanib. On the other hand, both TGF-β1 and TGF-β2 were lowered significantly by Nintedanib, but SB-431542 did not affect these levels as much, which is expected as SB-431542 acts more on TGFβ activation than on TGF-β expression levels (Figure 4f,i). Nintedanib and SB-431542 reduced MMP and TIMP protein expression levels closer to control healthy organoids (Figure 4g,j). Overall, our data show that the IPF-Fib with iAT2^I73T/tdT^ cell co-culture organoid model is amenable to drug testing and has the potential to identify the mechanisms of action of therapeutics.

## Discussion

There is a paucity of models that recapitulate the complex features of IPF, especially its initiation, progressive nature, tissue remodeling and loss of AT2 cells. Here, we present a cell co-culture, “top down” approach to model IPF that can be used to study different combinations of relevant cell types to derive a quantifiable readout of fibrosis. First, we showed that our cell co-culture organoid model develops cellular diversity, closely resembling that seen in IPF tissue. We saw the emergence of atypical basaloid cells, transitioning epithelial cells, and fibroblast cell subtypes, together with a general loss of epithelial cells while remodeling is occurring. Second, epithelial-fibroblast cell interactions induced secretion of fibrotic factors that we quantified and that corresponded with progression in the model. They were also used to assess the response to different compounds. Third, we used the model to assess the efficacy of therapies for IPF. We found a partial but dose dependent response to Nintedanib and the phosphodiesterase E4B inhibitor. This cell co-culture organoid model therefore has potential to be used to study the biology of IPF and perform drug discovery and testing. This model is complementary to another iAT2-iPSC-derived lung mesenchymal cell co-culture organoid model that also demonstrates fibrogenic cross-talk and cell changes with drug responses [43].

One of the unique features of our cell co-culture organoid model is the ability to combine cells from healthy and disease states. The epithelial-fibroblast cell combination of healthy fibroblasts with iAT2^tdT/WT^ cells showed phenotypic differences with the IPF fibroblasts with iAT2^I73T/tdT^ cells suggesting intrinsic cell differences that drive the fibrotic phenotype. However, by culturing iAT2^I73T/tdT^ cells with “healthy” fibroblasts in our cell co-culture organoid model we were able to mimic the initial effects of the I73T SFTPC mutation in AT2 cells on the surrounding fibroblasts. We presumed that this phenocopies the earliest changes induced in “healthy” fibroblasts by the iAT2^I73T/tdT^ cells. This particular cell co-culture combination induced the expression of genes known to respond to hypoxia. These include EGLN1 and EGLN3 that are part of a family of genes that are induced by hypoxia and act as an oxygen sensor that regulates the response to hypoxia-inducible factors, including upregulation of TGFβ [44, 45]. This suggests that this specific SFTPC mutation that results in protein misfolding and cell apoptosis could induce surrounding fibroblasts to sense hypoxia and respond accordingly. The model also allows the study of different lung fibroblast donors. We studied lung fibroblasts from three healthy donors and three IPF patients in the organoid models and we noticed heterogeneity in the production of COL1A1 and reduction in surface area of the organoid models. This is to be expected given variations in the population, and this could potentially be used to help tailor therapies for individuals.

Another unique feature of our cell co-culture organoid model is the ability to use scRNA sequencing to track cell subtypes over time. We especially noted the fibroblast cell subtypes with similarities to those that have been described in IPF tissue, including CTHRC1^+^/COL1A1^+^ fibroblasts that resemble fibrotic fibroblasts from fibroblastic foci, ACTA2^+^/COL1A1^+^ fibroblasts that resemble myofibroblasts, Prostaglandin expressing inflammatory fibroblasts, and fibroblasts with a specific cholesterol metabolism gene expression signature. Lipid metabolism in IPF is known to be dysregulated, and the model therefore provides a platform to study metabolic changes in different cell types during progressive fibrosis, and this could lead to interventions that could have therapeutic value. We also noted the appearance of K5^-^K17^+^ atypical basaloid cells over time in culture which is an epithelial cell type seen in IPF and some other fibrotic lung disorders. We found these cells arose not just from iAT2^I73T/tdT^ cells but also from AT2^tdT/WT^ cells. We speculate that this is because our cell co-culture model is developed on a stiff scaffold and that this stiffness in the microenvironment also induces AT2^tdT/WT^ cells to become atypical basaloid cells. Our model could, therefore, be used to screen for compounds that prevent the conversion of iAT2 cells into atypical basaloid cells, thereby reducing or halting progressive fibrosis.

Our cell co-culture organoid model also allows us to study the secreted proteins from the cell combinations and therefore this model could be particularly useful as a way of measuring biomarkers to determine how patients may respond to a particular drug, and ultimately as a marker of fibrotic disease burden. MMP7 is one of the five PROLIFIC biomarkers that has been found to be prognostic in the serum of IPF patients, and it was elevated in our IPF model and partially reduced with Nintedanib and SB-431542 treatment [46]. TIMP1 was also found to be elevated in the serum of patients with IPF as compared to healthy individuals and we found TIMP1 was elevated in our cell co-culture IPF model as compared to our cell co-culture healthy model and showed a partial response with Nintedanib but not with SB-431542. ECM proteins are also secreted by our model, and this alters the scaffold stiffness [47]. A unique feature of our model is that we can tune the scaffold to different stiffnesses and use this to assess cell responses to mechanotransduction signals over time. IPF is a progressive fibrotic disorder, and it is likely that cell-cell interactions change with alterations in the microenvironment. The ability to tune the microenvironment for IPF disease modeling is therefore essential for understanding progression of disease.

Given that our cell co-culture organoid model resembles the progressive fibrotic phenotype and cellular diversity seen in IPF, we tested Nintedanib and the Alk5 inhibitor SB-431542, to assess their efficacy in our model. The partial dose dependent response that we saw suggests that our IPF-Fib with iAT2^I73T/tdT^ cell co-culture organoid model could be used to assess the efficacy of therapeutics under development for the clinic and to predict clinical response. In addition, our model could also be used to test efficacy of combinations of drugs. This is particularly important, as many IPF clinical trials are enrolling patients who are still on Nintedanib or Pirfenidone.

## Conclusions

Herein, we present a scaffold-based human co-culture alveolar organoid model that recapitulates the progressive pathogenesis of pulmonary fibrosis. Combining induced pluripotent stem cell-derived alveolar type 2 (iAT2) cells and primary lung fibroblasts, this model elucidates critical signaling axes underlying fibrosis initiation and progression, including epithelial-mesenchymal crosstalk, extracellular matrix remodeling, and dysregulated cytokine and chemokine secretion. Furthermore, the model demonstrates dose-dependent responsiveness to clinically approved anti-fibrotic therapeutics, with half-maximal inhibitory concentration (IC50) values concordant with those observed in clinical settings. Overall, our cell co-culture organoid model of progressive fibrosis highlights the importance of cell-cell interactions and the microenvironment in IPF and how more complex organoid models could help with future drug discovery and guide the advancement of drug candidates to clinical trials.

## Limitations of our study

Models, by their very nature, are a reductionist approach to mimic disease. Our IPF model is particularly useful for studying cell-cell interactions in the earliest phase of the disease and following progression over time. It is, however, not truly representative of the Usual Interstitial Pneumonia (UIP) histopathology, and other major cell types that play a role in IPF are not included in this iteration of the model. For example, our model lacks alveolar macrophages that are patterned during development and play a critical role as a first line of host defense but are also critical for SFTP turnover and are likely important in IPF initiation and progression. However, our cell co-culture organoid model is amenable to integration of other cell types due to its “top down” approach. Our model also lacks vascularization with endothelial cells and blood flow, which in turn can affect inflammatory cell influx and redox studies. Newer vascularization platforms are now available, and these can be incorporated into the model to further refine it.

## Resource availability

### Lead contact

Further information and requests for resources and reagents should be directed to and will be fulfilled by the Lead Contact, Brigitte N. Gomperts.

## Acknowledgments

This work was supported by National Heart, Lung, and Blood Institute (NHLBI) U01HL153000 (to B.N.G.), NHLBI LungMAP U01HL175451 (to B.N.G.), DoD PR202868 (to B.N.G.) and the Burroughs Wellcome Fund 1020030 (to B.N.G.). B.N.G. was also supported by the UCLA BSCRC and the Jonsson Comprehensive Cancer Center. This work was also supported by NIH grants R01HL095993, NO175N92025D00035 and U01HL148692 to DNK, NIH K08 HL163494 and a Boston University School of Medicine Department of Medicine Career Investment Award to KDA, NIH P01HL170952 and U01HL152976, and a research grant from GSK to DNK and KDA, and Swiss National Science Foundation P500PB_206631 to ABA. We appreciate the UCLA BSCRC Microscopy Core, the UCLA Translational Pathology Core Laboratory (TPCL), and the UCLA Clinical and Translational Science Institute (CTSI).

## Author contributions

Conceptualization, C.S and B.N.G.; data curation, C.S. and J.L.; formal analysis, C.S and J.L., funding acquisition, B.N.G.; investigation, C.S., B.N.G., K.P., D.N.K.; experimentation, C.S., J.L., K.A., A.B.A., C.C., K.C., S.S., R.R.C., W.C., T.R.; methodology, C.S., J.L., K.P., and B.N.G.; software, J.L; supervision, K.P., and B.N.G., writing – original draft, C.S.; writing – review & editing, K.P. and B.N.G.

## Supplemental Information

Supplemental figures are in a separate pdf.

## Ethical Approval and consent to participate

All patient samples used in this study were de-identified and were collected with an Institutional Review Board (IRB) exemption. The need for consent was waived by the IRB. Patient large airways and bronchial tissues were acquired from three different tissue sources: 1. de-identified normal human donors after lung transplantations at the Ronald Reagan UCLA Medical Center. Tissues were procured under an Institutional Review Board-approved protocol at the David Geffen School of Medicine at UCLA (IRB#21-000390-CR-00003). 2. Normal human bronchial epithelial cells (NHBE) from non-smokers were obtained from Lonza and all samples were de-identified. 3. Deidentified donor lung and trachea samples from the International Institute for the Advancement of Medicine (IIAM) were obtained with institutional approval. All animal studies were carried out according to the standard operating procedure in place at the test facility at Vibiosphen. All procedures were performed in accordance with the Directive 2010/63/UE recommendations and with French Veterinary Authorities agreement. All animals were managed similarly, with due regard for their well-being, according to prevailing practices and the current SOPs in force at Vibiosphen. The in vivo design and procedures were approved by Ethical Committee.

