## Supplemental Figures and legends for "Modeling cell-cell interactions to advance drug discovery in Idiopathic Pulmonary Fibrosis"

### Supplemental Figure 1

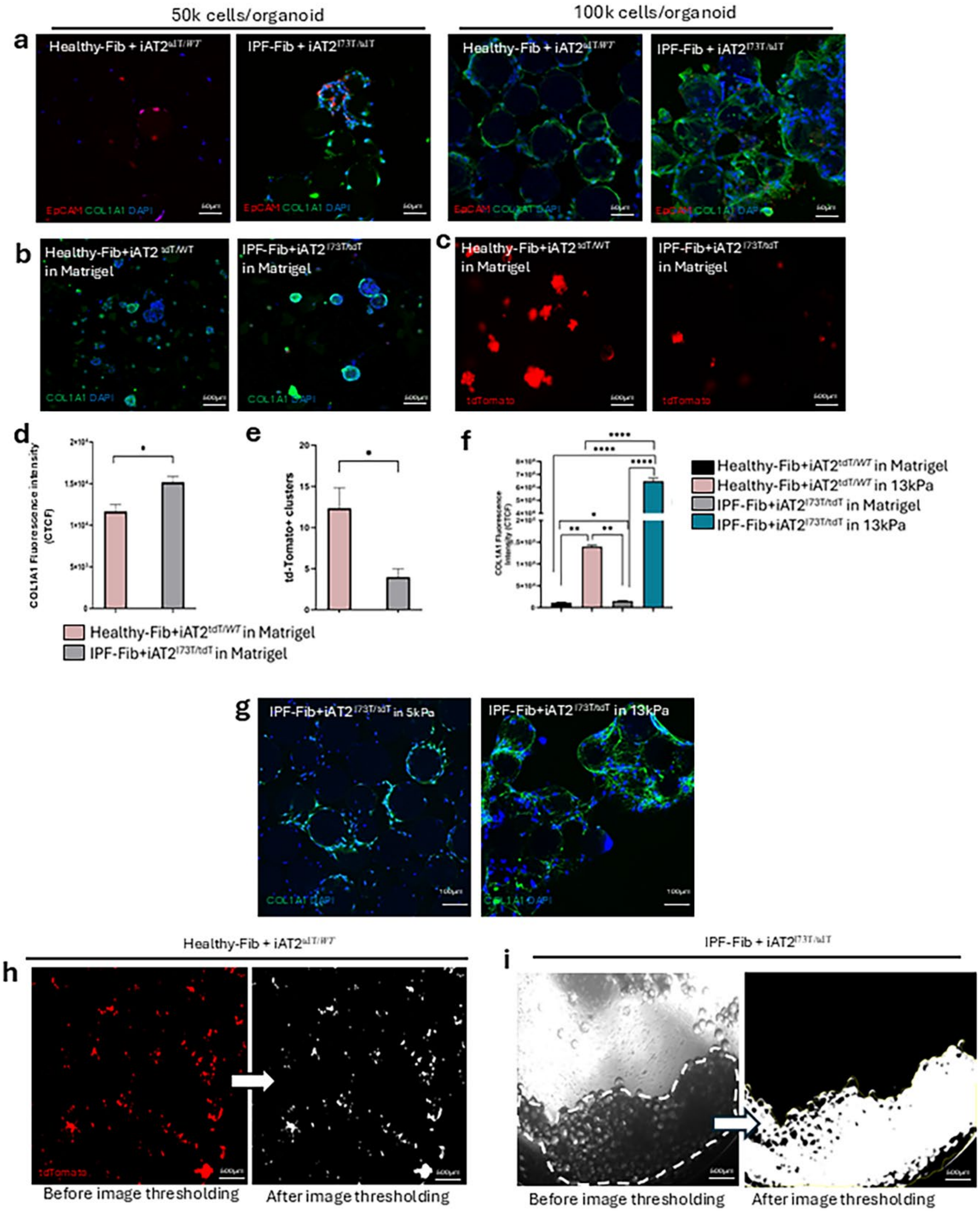

#### Supplemental Figure 2

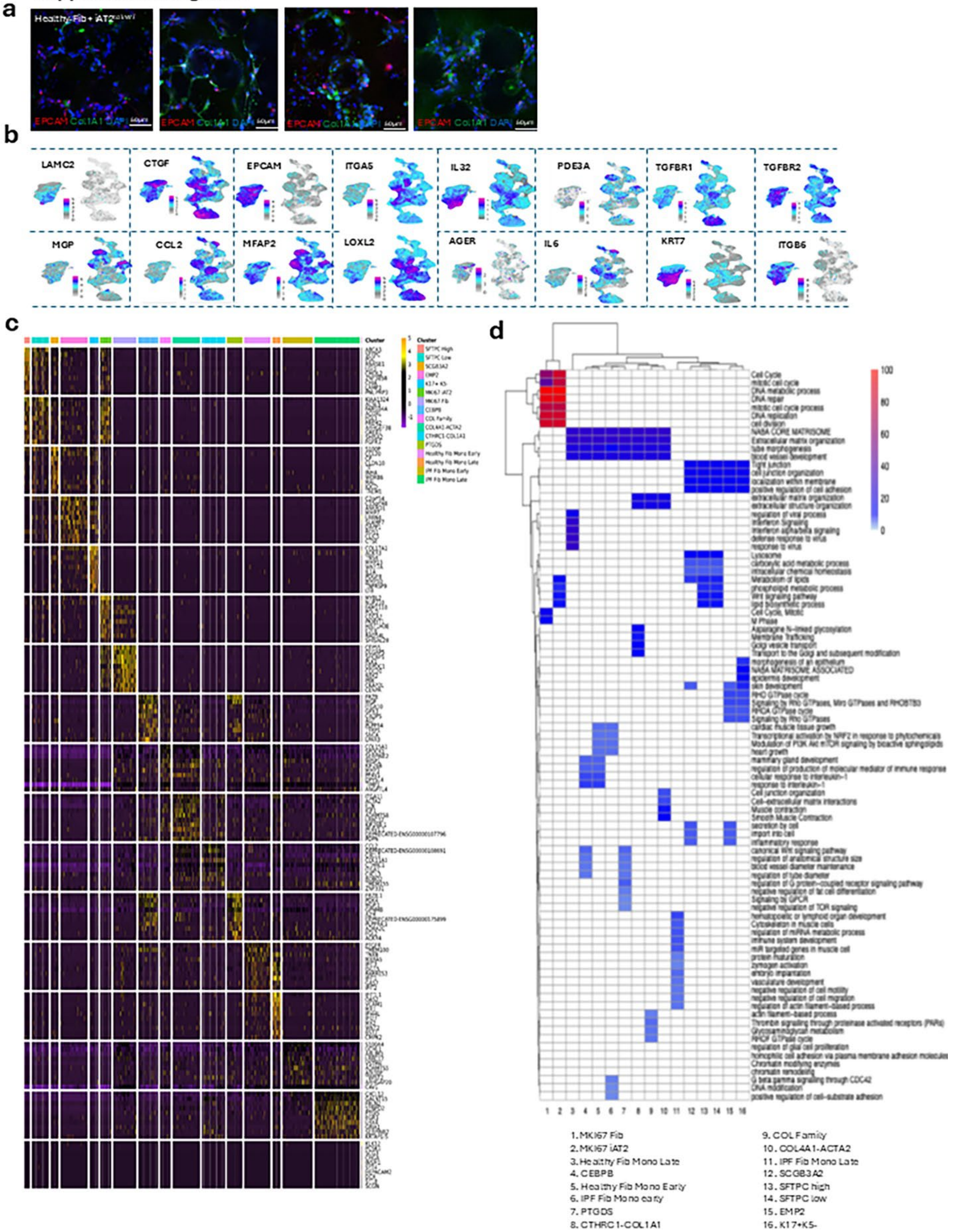

**Supplemental Figure 3**

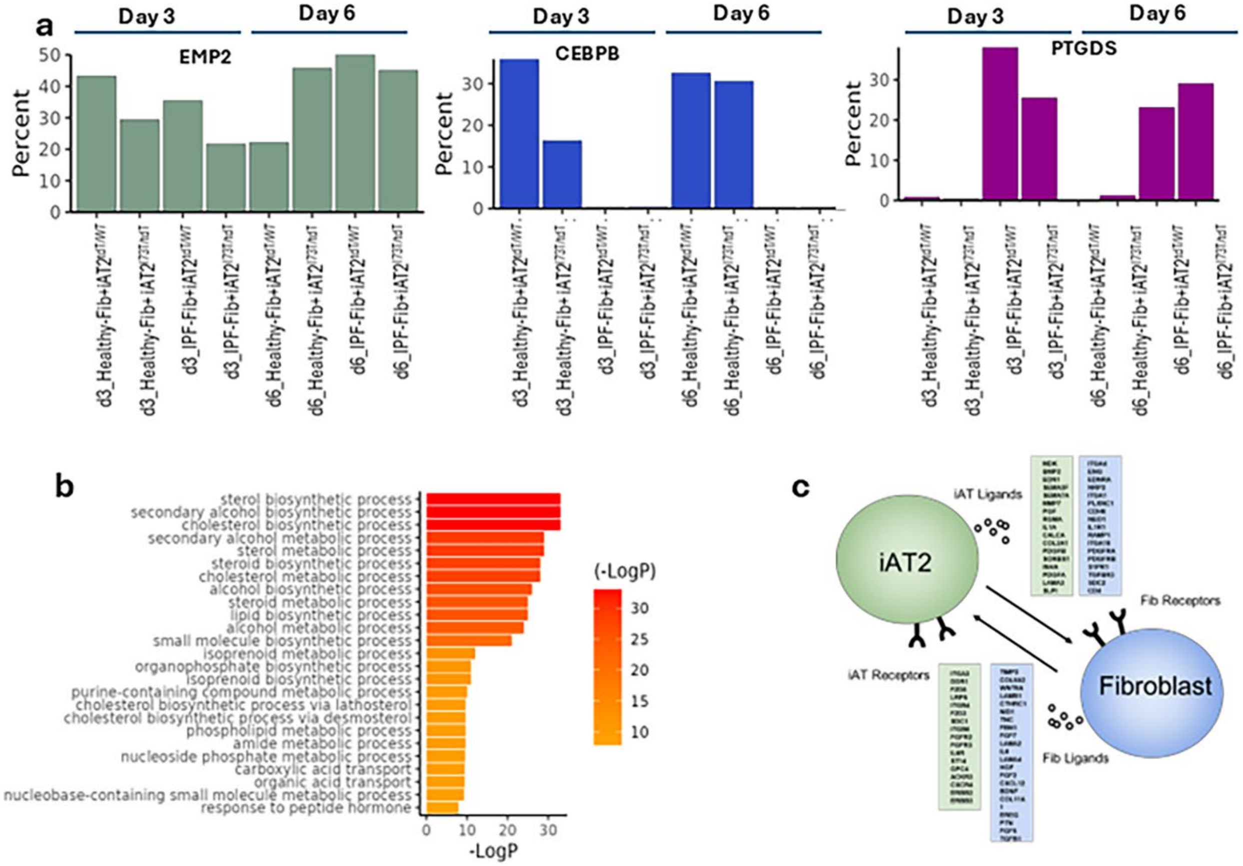

**Supplemental Figure 4**

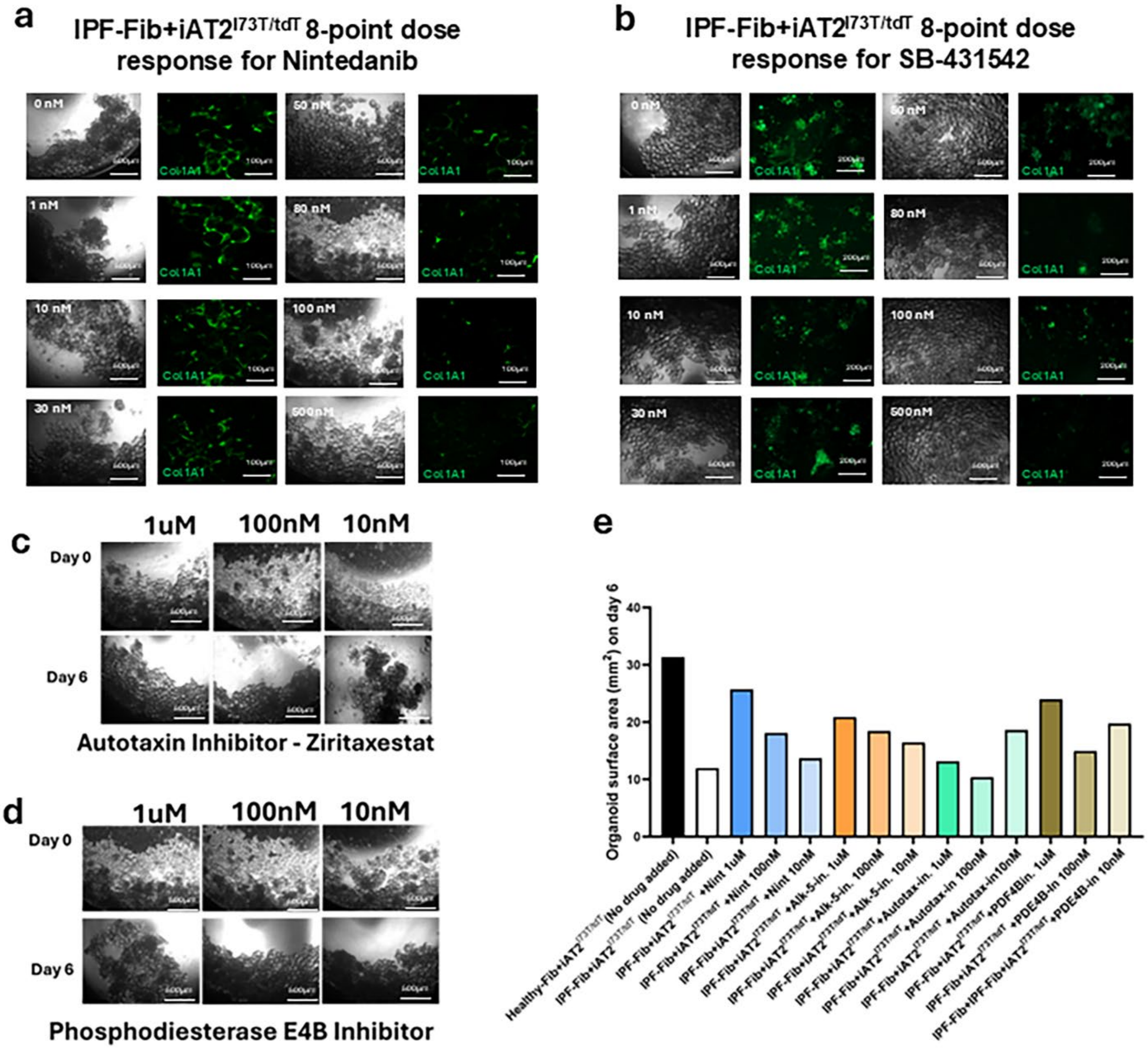

#### Supplementary Figure Legends

##### Supplementary Figure 1. Model optimization:

**a.** Effect of cell numbers (50,00 cells per organoid vs. 100,000 cells per organoid) in the model formation and corresponding EpCAM and COL1A1 expressions by immunofluorescent staining. Scale bar = 50µm. **b-c.** Co-culture of Healthy-Fib+iAT2$^{tdT/\mathit{WT}}$ and IPF-Fib+iAT2$^{I73T/tdT}$ cells in the Matrigel droplets on day 6: **b.** COL1A1 expression and **c.** tdTomato$^{+}$ cells. Scale bar = 500µm. **d-e.** Quantification of COL1A1 fluorescence intensity and tdTomato$^{+}$ cells in Matrigel droplets. Student's t-test; *= P<0.05. **f.** Quantification of COL1A1 fluorescence intensity of Healthy-Fib + iAT2$^{tdT/\mathit{WT}}$ cell co-culture and IPF fibroblasts + iAT2$^{I73T/tdT}$ cell co-culture in Matrigel droplets and in 13kPa microbeads on day 6. One-way ANOVA; *= P<0.05, **= P<0.01, ****= P<0.0001. **g.** COL1A1 expression in fibroblasts seeded on 5kPa microbeads versus 13kPa microbeads on day 6. Scale bar = 100µm. **h-i.** The accuracy of image segmentation is determined by cell number count and surface area quantification (see image analysis in methods). Scale bar = 500µm.

##### Supplementary Figure 2. Specific genes and gene networks in clusters:

**a.** Immunofluorescence images (Day 3) of EpCAM and COL1A1 expression in the co-culture combinations (Healthy-Fib+iAT2$^{tdT/\mathit{WT}}$,Healthy-Fib+iAT2$^{I73T/tdT}$,IPF-Fib+iAT2$^{tdT/\mathit{WT}}$,IPF-Fib+iAT2$^{I73T/tdT}$) used in sc-RNA seq experiment. Scale bar = 50µm. **b.** UMAPs of specific genes expressed in the cell clusters. **c.** Heatmap of top 10 differentially expressed genes (DEGs) of each cluster. **d.** Gene Ontology (GO) analysis of enriched gene networks in each cluster.

##### Supplementary Figure 3. Changes in clusters with time:

**a.** Changes in proportions of EMP2, CEBPB, and PTGDS clusters per sample on day 3 and day 6 of culture. **b.** GO terms related to lipid metabolism. **c.** Identified receptor-ligand interactions in the model.

##### Supplementary Figure 4. Testing of efficacy of clinical compounds in the IPF-Fib with iAT2$^{I73T/tdT}$ cell co-culture organoid model.

**a.** IPF-Fib with iAT2$^{I73T/tdT}$ cell co-culture organoid model 8-point dose course response to Nintedanib with brightfield images (Scale bar = 500µm) and IF (Scale bar = 100µm) for Col1A1 expression in green. **b.** IPF-Fib with iAT2$^{I73T/tdT}$ cell co-culture organoid model 8-point dose course response to the SB-431542 (Alk5 inhibitor) with brightfield images (Scale bar = 500µm) and IF (Scale bar = 100µm) for Col1A1 expression in green. **c.** Brightfield microscopic images of IPF-Fib with iAT2$^{I73T/tdT}$ cell co-culture organoid model showing model surface area on days 0 and 6 (Scale bar = 500µm) treated with 1uM, 100nM, and 10nM of autotaxin inhibitor Ziritaxestat. **d.** Brightfield microscopic images of IPF-Fib with iAT2$^{I73T/tdT}$ cell co-culture organoid model showing model surface area on days 0 and 6 (Scale bar = 500µm) treated with 1uM, 100nM, and 10nM of phosphodiesterase E4B inhibitor (Nerandomilast) and **e.** Comparison of organoid surface area on day 6: from left to right- no drug added Healthy-Fib with iAT2$^{tdT/\mathit{WT}}$ cell co-culture organoid model, no drug added IPF-Fib with iAT2$^{I73T/tdT}$ cell co-culture organoid model, and IPF-Fib with iAT2$^{I73T/tdT}$ cell co-culture organoid model treated separately with Nintedanib, Alk-5 inhibitor (SB-431542), autotaxin-inhibitor (ziritaxestat) and PDE4B inhibitor (Nerandomilast). All drugs were added in three concentrations- 1μM, 100nM and 10 nM.
